# Epigenetic signals that direct cell type specific interferon beta response in mouse cells

**DOI:** 10.1101/2022.02.26.481127

**Authors:** Markus Muckenhuber, Isabelle Lander, Katharina Müller-Ott, Jan-Philipp Mallm, Lara C. Klett, Caroline Knotz, Jana Hechler, Nick Kepper, Fabian Erdel, Karsten Rippe

## Abstract

The antiviral response induced by type I interferon (IFN) via the JAK-STAT signaling cascade activates hundreds of IFN-stimulated genes (ISGs). While this response occurs essentially in all human and mouse tissues it varies between different cell types. However, the linkage between the underlying epigenetic features and the ISG pattern of a given cell is not well understood. We mapped ISGs, binding sites of the STAT1 and STAT2 transcription factors and chromatin features in three different mouse cell types (embryonic stem cells, neural progenitor cells and embryonic fibroblasts) before and after treatment with IFNβ. The analysis included gene expression, chromatin accessibility and histone H3 lysine modification by acetylation (ac) and mono-/tri-methylation (me1, me3). A large fraction of ISGs and STAT binding sites were cell type specific with promoter binding of a STAT1-STAT2 complex (STAT1/2) being a key driver of ISG induction. Furthermore, STAT1/2 binding to putative enhancers at intergenic and intronic sites induced ISG expression as inferred from a chromatin co-accessibility analysis. STAT1/2 binding was dependent on the chromatin context and positively correlated with pre-existing H3K4me1 and H3K27ac marks in an open chromatin state while the presence of H3K27me3 had an inhibitory effect. Thus, chromatin features present before stimulation represent an additional regulatory layer for the cell type specific antiviral response.

## Introduction

Type I interferons (IFNs) like IFNα and IFNβ are expressed across almost all tissues in human and mouse as a first line of defense against viral infections (Hoffmann et al, 2015; Lazear et al, 2019; Sa Ribero et al, 2020; Stanifer et al, 2020). They activate hundreds of IFN-stimulated genes (ISGs) during innate immune response. Virus infection induces IFNβ in most cell types, which then can stimulate production of other type I IFNs (Hoffmann et al, 2015). The ISG activation by IFN is not uniform but occurs in a cell type specific manner (Lazear et al, 2019; Sa Ribero et al, 2020; Stanifer et al, 2020) and displays striking changes during differentiation of human embryonic stem cells (Wu et al, 2018). Mouse embryonic stem cells (ESCs) do not express IFN themselves upon viral infection but respond to IFN and display an attenuated innate immune response as compared to differentiated murine cells (D’Angelo et al, 2016; Gonzalez-Navajas et al, 2012; Guo, 2017; Guo et al, 2015; Wang et al, 2014; Wang et al, 2013; Whyatt et al, 1993).

One aspect of the cell type specific response to IFNs are specific epigenetic features that modulate ISG activation via the JAK-STAT signaling cascade. This pathway involves phosphorylation of STAT1 and STAT2 transcription factors that, together with IRF9, assemble into the IFN-stimulated gene factor 3 (ISGF3) complex (Chen et al, 2017; Hu et al, 2021; Ivashkiv & Donlin, 2014; Stark & Darnell, 2012; Villarino et al, 2017). ISGF3 translocates into the nucleus, binds interferon-stimulated response elements (ISREs) and activates ISGs. In addition, IFNγ activation sites (GAS) are bound predominantly by phosphorylated STAT1 homodimers and can drive IFN mediated gene induction. The STAT binding sites are frequently located at promoters and regulatory sites such as enhancers (Begitt et al, 2014; Ostuni et al, 2013; Vahedi et al, 2012). Chromatin remodeling complexes, histone acetyltransferases and deacetylases can act as modulators for the downstream JAK-STAT signaling cascade (Au-Yeung & Horvath, 2018; Chen et al, 2017; Liu et al, 2002; Nusinzon & Horvath, 2003; Testoni et al, 2011; Villarino et al, 2017). However, it is not well understood how specific chromatin features affect STAT1 and STAT2 binding and ISG induction. Here, we dissected the cell type specific IFNβ response by comparing mouse ESCs, neural progenitor cells (NPCs) derived by *in vitro* ESC differentiation and mouse embryonic fibroblasts (MEFs) in a comprehensive genome-wide analysis. Our sequencing-based readouts comprised transcription, binding of STAT1 and STAT2, acetylation (ac) and mono- and tri-methylation (me1, me3) of histone H3 lysine residues (H3K4me1, H3K4me3, H3K9ac, H3K27ac, H3K9me3 and H3K27me3) and open chromatin mapped by the assay for transposase-accessible chromatin (ATAC). The resulting sets of common and cell type specific ISGs were linked to the binding of a STAT1-STAT2 complex (STAT1/2) at promoters and enhancers in dependence of their chromatin state. Our analysis sheds light on the interplay of epigenetic signals, STAT1/2 binding at cis-regulatory elements and the cell type specific modulation of innate immune response.

## Results

### IFNβ induces anti-viral gene expression programs in all three cell types

ESCs, MEFs and NPCs were obtained from a 129/Ola mouse strain and represent an established cellular system that allowed us to compare the cell type dependent epigenetic makeup and IFNβ response of the same genome for a large number of chromatin features (Molitor et al, 2017; Teif et al, 2012) (**Fig. 1A**). The three different cell types were treated with IFNβ for 1 h or 6 h and gene expression profiles were acquired by RNA sequencing (RNA-seq) (**Supplementary Table S1**). Differential gene expression analysis identified in total 191 ISGs induced in ESCs, 463 ISGs in MEFs, and 244 ISGs in NPCs over unstimulated controls (0 h) (**Fig. 1B, Supplementary Table S2, Supplementary Data Set 1**). As expected, a GO-term analysis yielded upregulated genes related to anti-viral programs and innate immune responses in all three cell types (**Supplementary Fig. 1A**). By intersecting the three individual ISG sets, we obtained 143 common ISGs while 33 (ESC), 17 (NPC) and 221 (MEF) ISGs were cell type specific (**Fig. 1C, Supplementary Data Set 1**). The ISGs found in NPCs mainly represented a subset of MEF ISGs (227 of 244) pointing to a high similarity of the IFNβ response in NPCs and MEFs (**Fig. 1C**). A differential gene expression analysis of only intronic reads to assess nascent RNA levels gave very similar results with a somewhat lower number of ISGs detected in ESCs (**Supplementary Fig. 1B, C; Supplementary Table S2**). We conclude that changes induced by IFNβ occurred predominantly at the gene expression level with only minor differences in RNA stability.

**Figure 1.**
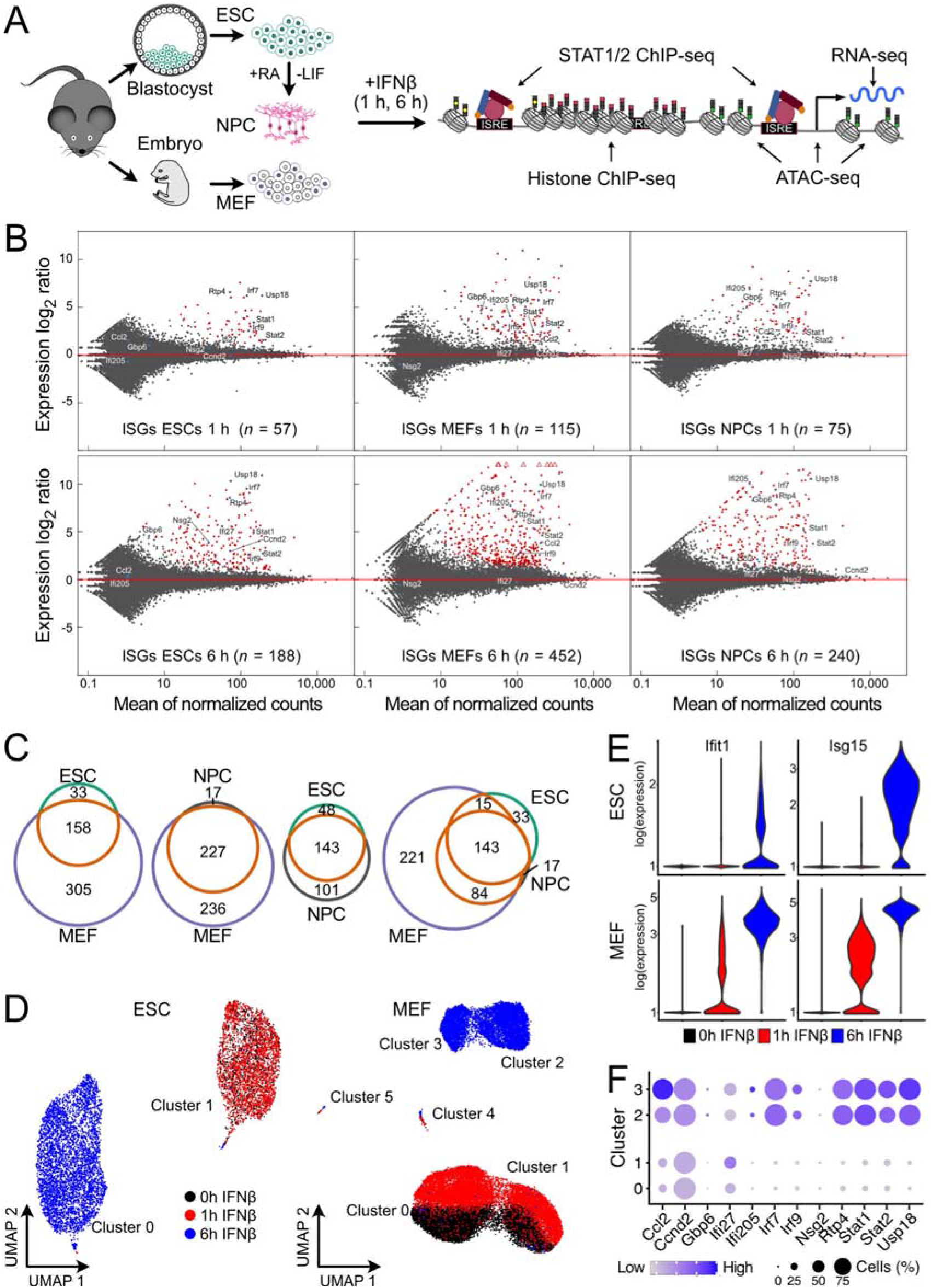
ISG induction patterns in ESCs, MEFs and NPCs. (**A**) ESCs, NPCs differentiated *in vitro* from them and MEFs from a 129/Ola mouse strain were studied to reveal the relation between cell type specific chromatin features and IFNβ response. (**B**) Gene expression changes after IFNβ treatment. Red dots represent significant differentially expressed genes at *p*_adj_ < 0.05 and fold change ≥ 1.5. Four biological replicates for ESCs, two for MEFs, and four for NPCs were acquired for RNA-seq. (**C**) Overlap of ISGs found after 1 h or 6 h IFNβ treatment in ESCs, NPCs and MEFs. (**D**) Singlecell embedding of gene expression in ESCs (left) and MEFs (right). (**E**) Normalized expression levels of the ISGs *Ifit1* and *Isg15* in single ESCs (top) and MEFs (bottom). Both genes were reliably detected as ISGs in the bulk RNA-seq analysis after 1 h. (**F**) Expression levels of selected ISGs identified by bulk RNA-seq data according to aggregated scRNA-seq in MEF clusters 0, 1, 2 and 3.

### IFNβ response is mostly homogenous at the single-cell level

We assessed by single-cell RNA sequencing (scRNA-seq) if the transcriptional response in ESCs and MEFs was homogeneous or if the observed upregulation of ISGs arises from a subset of strongly responding cells (**Fig. 1D; Supplementary Fig. 1D**). In contrast to ESCs, MEFs consistently formed two distinct clusters (clusters 0 and 1 and clusters 2 and 3, respectively) in the single-cell embedding of transcriptomic profiles. This clustering arose from upregulated genes associated with KEGG pathway “extra cellular matrix receptor interaction” in clusters 0 and 2 as opposed to the “focal adhesion” KEGG pathway in clusters 1 and 3 (**Supplementary Fig. 1E, F**). Based on these expression profiles we annotated clusters 0 and 2 as “mesenchymal-like” and clusters 1 and 3 as “epithelial-like”.

Inspection of the UMAP plots showed no separate clustering of untreated (0 h) and 1 h IFNβ treated ESCs, while cells after 6 h stimulation formed a distinct cluster. Untreated and 1 h IFNβ treated MEFs separated within the same clusters while MEFs treated for 6 h with IFNβ were present in a separate cluster again. The pattern was in line with the relatively small global transcriptomic changes in ESCs and MEFs after 1 h IFNβ treatment, where 57 and 115 ISGs were detected by bulk RNA-seq as compared to 188 and 452 genes after 6 h for ESCs and MEFs, respectively (**Fig. 1B**). The scRNA-seq response pattern was illustrated for two ISGs, *Ifit1* and *Isg15* (**Fig. 1E**). The number of cells where transcripts of the two genes were detected, largely increased from the 1 h to the 6 h time point as more RNA is produced. We conclude that the apparent heterogeneity after 1 h IFNβ appears to arise to a significant extend from the reduced detection sensitivity of scRNA-seq for lowly expressed genes that show an increased drop-out frequency (Yamawaki et al, 2021). Furthermore, the expression patterns and IFNβ response dynamics of the two MEF clusters (cluster 0 vs 1; cluster 2 vs 3) were highly similar in terms of ISGs and their induced gene expression levels (**Fig. 1F**). Thus, the IFNβ response was rather homogeneous in the two different cell types at the single-cell level and we used the ISG definition from the bulk RNA-seq analysis for further analysis.

### ISG expression varies between cell types in response strength and specificity

Next, we compared the transcriptional response to IFNβ in the three cell types in further detail. The distribution of gene expression levels in non-stimulated cells was fitted with distributions for active and repressed genes to define a background threshold for evaluation of differences in the IFNβ response (**Supplementary Fig. 2A**). In ESCs and MEFs some genes like *Irf9, Stat1* and *Stat2* were already lowly expressed in unstimulated cells and showed a significant increase in expression after IFNβ treatment (**Fig. 2A**). Other ISGs like *Irf7, Rtp4* and *Usp18* changed from a repressed to an activated state after IFNβ stimulation. Compared to ESCs, MEFs displayed a 10 to 100-fold stronger induction of these common ISGs, which is in line with previous findings (Wang et al, 2014). To further dissect the overall stronger response in MEFs, we compared expression levels of factors of the IFN signaling pathway. The *Ifnar1* and *Ifnar2* receptors as well as *Jak1* kinase were higher expressed in MEFs than in ESCs while for key transcription factors *Stat1, Stat2* and *Irf9* no differences were identified (**Supplementary Fig. 2B**). A western blot with STAT1 and STAT2 antibodies showed that STAT1 and STAT2 proteins were present at lower levels in ESCs before and after IFNβ induction as compared to MEFs (**Fig. 2B**). The amount of STAT1 phosphorylated at residue 701 (STAT1_p701_) or 727 (STAT1_p721_) was clearly increased after 1 h in MEFs as compared to ESCs and decayed to low levels at the 6 h time point. Thus, we conclude that the globally attenuated response to IFNβ in ESCs involved epigenetic networks that lead to a reduced expression of key components of the JAK/STAT signaling pathway as compared to differentiated cells. The lower levels of active STAT1/2 protein complexes upon IFNβ induction are apparent from the comparing the amounts of STAT1_p701_ and STAT1_p727_ between ESCs and MEFs. In addition to these global differences, cell type specific differences were apparent as illustrated for selected genes in **Fig. 2C**. After 6 h of stimulation *Ccnd2, Ifi27*, and *Nsg2* were induced in ESCs. In NPCs, *Ccnd2* and *Nsg2* were constitutively expressed while the lowly expressed *Ifi27* showed only a small expression increase after 6 h. In MEFs, expression of all three genes was not upregulated. In contrast, *Ccl2, Gbp6* and *Ifit1bl1* were specifically upregulated in MEFs upon IFNβ stimulation. *Gpb6* and *Ifit1bl1* also showed a response in NPCs albeit at a lower level. *Gbp6* was lowly induced in ESCs but only after 6 h. In summary, large cell type specific differences in gene expression levels were observed upon IFNβ stimulation between the three cell types that involved the expression of distinct sets of ISGs in ESCs and MEFs with NPCs showing a pattern that was similar to MEFs.

**Figure 2.**
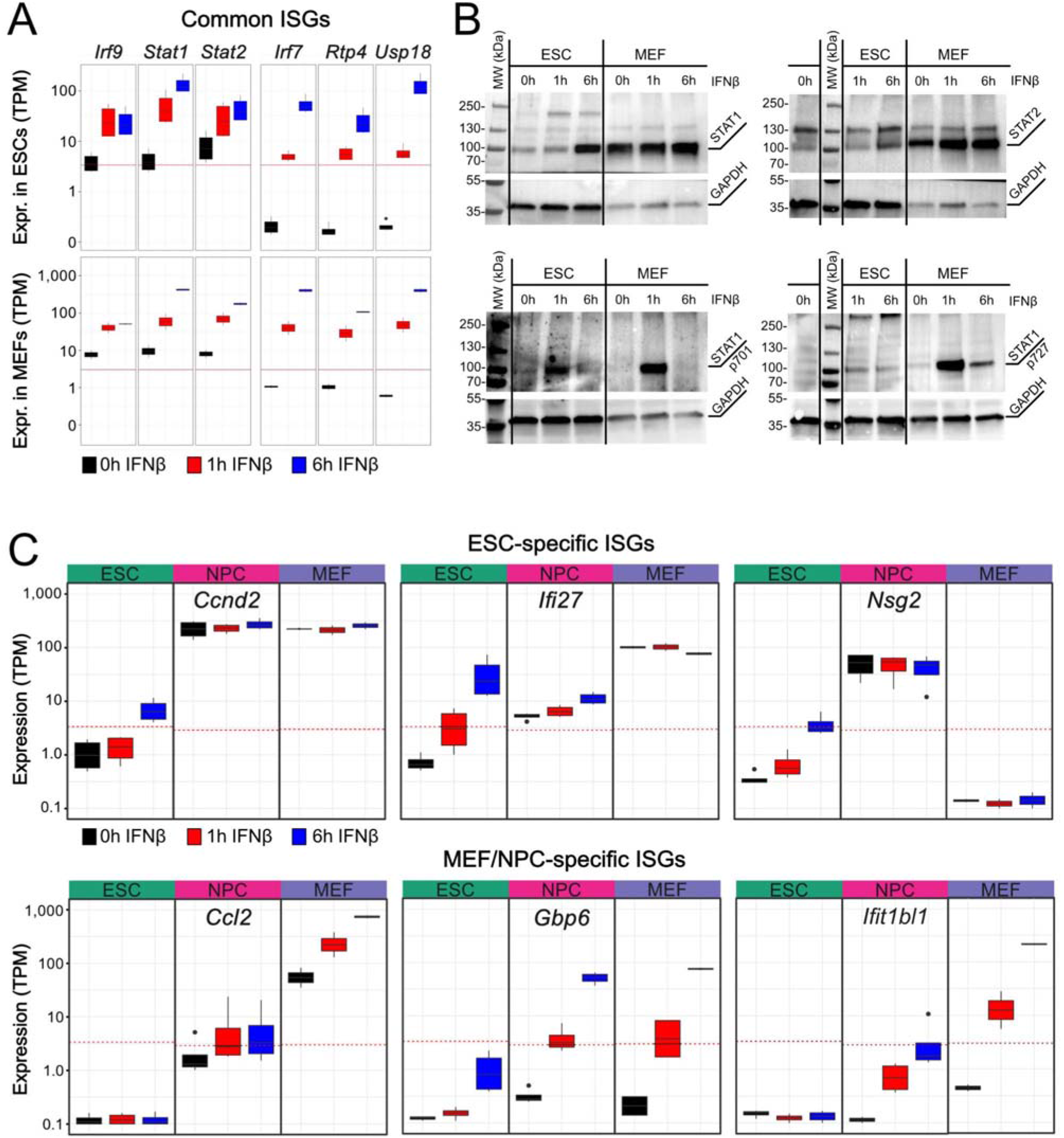
Cell type specific ISG induction and protein expression. (**A**) Normalized gene expression levels of selected ISGs from bulk RNA-seq in ESCs (top) and MEFs (bottom). Gene expression is given as transcripts per kilobase million (TPM). (**B**) Western blots of IFNβ stimulated ESCs and MEFs at 0 h, 1 h and 6 h time points. The top row shows total levels of STAT1 (left) and STAT2 (right). The lower row shows phosphorylation of STAT1 at position 701 (left) and 727 (right). GAPDH was used as housekeeper gene control. (**C**) Normalized gene expression levels of selected cell type-specific ISGs in ESCs, NPCs and MEFs. The red line represents a cell type-specific threshold to distinguish active and repressed genes. Top: Expression of ISGs *Ccnd2, Ifi27* and *Nsg2* was only induced in ESCs. Bottom: Expression of ISG *Ccl2* was induced in MEFs. Expression of ISGs *Gbp6* and *Ifit1bl1* was induced in MEFs and NPCs.

### STAT1/2 binding is cell type specific and correlates with ISG activation

The differences in IFNβ response raise the question why certain ISGs were preferably expressed in one cell type and not in the other. To reveal molecular details of gene expression regulation we mapped STAT1_p701_ and STAT2 binding by chromatin immunoprecipitation followed by sequencing (ChIP-seq). Antibodies against STAT1_p701_ and STAT2 in ESCs and MEFs were used with exemplary regions enriched for both transcription factors shown in **Fig. 3A** and the number of peaks detected given in **Supplementary Table S2**. A total of 208 peaks in ESCs and 276 peaks in MEFs were bound simultaneously by both transcription factors (**Fig. 3B, Supplementary Table S2, Supplementary Data Set 2**). These loci were annotated as “STAT1/2” binding sites in our analysis. They were likely to represent the ISGF3 complex as it has been shown previously, that STAT1 and STAT2 assemble with IRF9 to form the ISGF3 complex upon IFN stimulation (Platanitis et al, 2019). A total of 392 STAT1/2 binding sites were determined from the combined data set of ESCs and MEFs after 1 h and 6 h of IFNβ stimulation. The remaining peaks that only had STAT1_p701_ or STAT2 bound were classified as “STAT1” and “STAT2” binding sites, respectively. The overlap of peaks between cell types was moderate (**Fig. 3E**). Only 38 sites were found to be bound by STAT1 in both cell types, while most STAT2 peaks were cell type specific. STAT1/2 binding sites common to both cell types comprised 44% (ESC) and 33% (MEFs) of the peaks. To validate the peak specificity, we determined enriched known motifs in STAT binding sites. In both ESCs and MEFs, the STAT-family motifs (STAT1, STAT3, STAT3 + IL21, STAT4, STAT5) were enriched at STAT1 peaks, while IRF-family motifs (IRF1, IRF2, IRF3, IRF8, ISRE) were the most enriched motifs in STAT1/2 and STAT2 peaks (**Fig. 3C**). Due to the high similarity of motifs within each family, we defined all detected STAT peaks with at least one of these to be a specific peak (**Supplementary Fig. 3A**). At least one of these family motifs was found in 66% (STAT1), 83% (STAT1/2) and 86% (STAT2) of the ESC peaks and 85% (STAT1), 90% (STAT1/2) and 88% (STAT2) of the MEF peaks. Thus, the same motifs were recognized independent of cell type and in line with the classification into STAT1, STAT2 and STAT1/2 binding sites. This conclusion was corroborated by a de novo motif analysis (**Supplementary Fig. 3B-C**). The top de novo motif was in all groups one of the STAT or IRF family with a similarity score of ~0.9. It is noted that the total number of the 1,885 STAT peaks detected by ChIP-seq represents only a minor fraction of the approximately 2.5 million STAT-or IRF-family sequence motifs in the mouse genome (~0.8 million IRF motifs, 1.7 million STAT motifs). Based on these findings, we conclude that the DNA sequence is neither sufficient to predict the experimentally observed STAT binding sites nor can it rationalize the differences in STAT binding sites detected between cell types.

**Figure 3.**
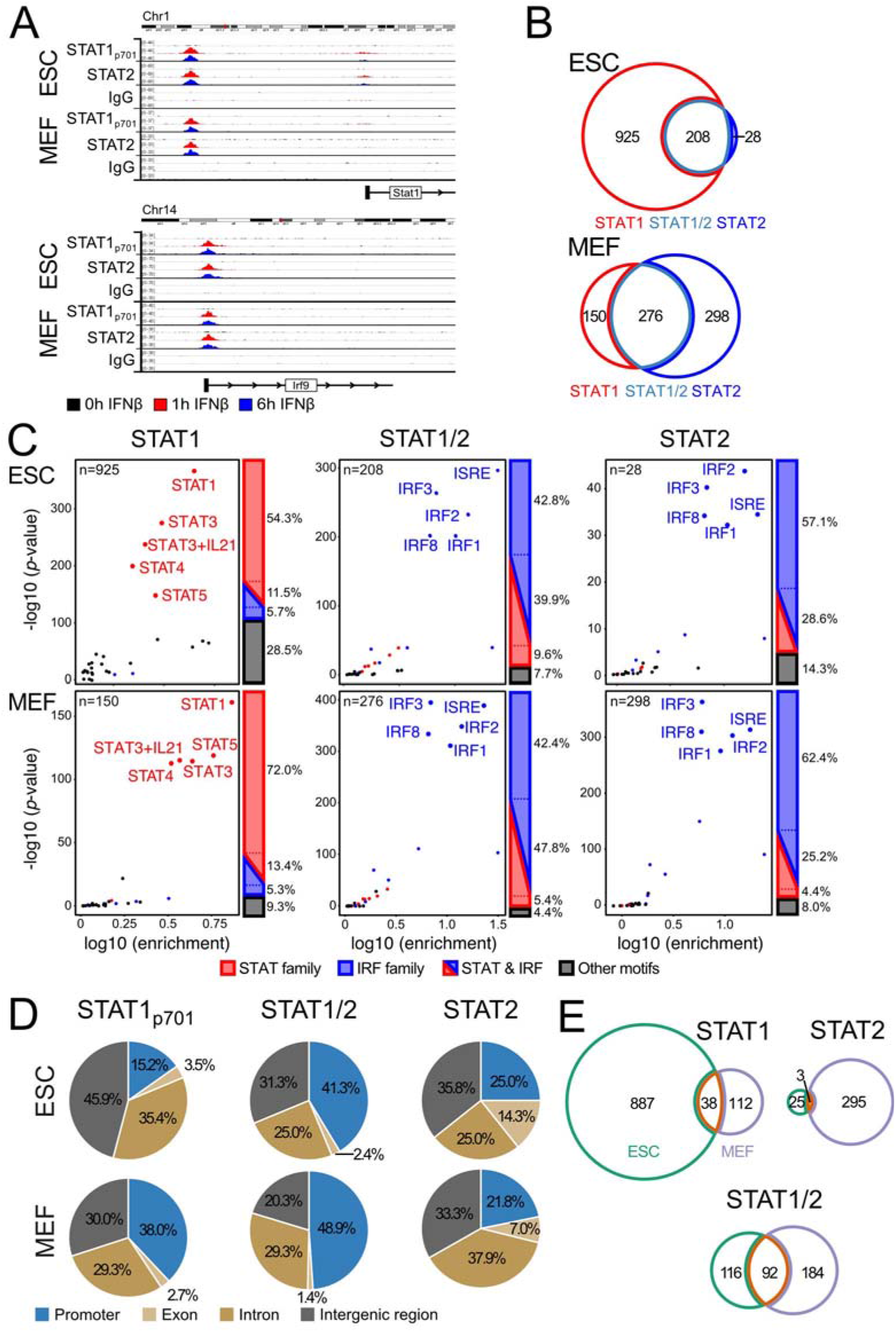
Cell type specific binding of STAT1 and STAT2. (**A**) ChIP-seq of STAT1_p701_ and STAT2 in the genomic regions upstream of *Stat1* (left) and at the promoter of *Irf9* (right). Tracks show one representative replicate for each condition. (**B**) STAT1_p701_ and STAT2 peaks in ESCs and MEFs. The overlap of STAT1_p701_ and STAT2 peaks defined STAT1/2 binding sites. (**C**) Enrichment of transcription factor binding motifs in STAT1_p701_, STAT1/2 and STAT2 peak sets identified in ESCs and MEFs. Motif color scheme: STAT-family (STAT1, STAT3, STAT3+IL21, STAT4, STAT5), red; IRF-family (IRF1, IRF2, IRF3, IRF8 and ISRE (IRF9)), blue; other, black. Four biological replicates for ESCs and two for MEFs were analyzed. (**D**) Distribution of STAT1_p701_, STAT1/2 and STAT2 peaks at promoters, exons, introns and intergenic regions annotated from the ENSEMBL data base. (**E**) Overlap of STAT binding sites between ESCs and MEFs for STAT1_p701_, STAT1/2 and STAT2.

### ISG activation can be partly assigned to STAT promoter binding

To gain further insight in the activation mechanism of ISGs, we analyzed the spatial relation between STAT binding sites and ISGs. Almost half of the STAT1/2 peaks in ESCs and MEFs were located at promoters (defined as a window of ± 1 kb around the transcription start site) with around 3/4 of them at ISGs (**Fig. 3D, Supplementary Fig. 3D, Supplementary Data Set 3**). In contrast, a smaller fraction of 15-38% of the STAT1 or STAT2 only peaks were at promoters. In addition, the promoters that displayed STAT1 binding but lacked STAT2 were mostly highly expressed genes and only a minor fraction of 6% in ESCs and 16% in MEFs were at ISGs. This fraction was around 50% for the STAT2 only peaks. Based on this analysis we conclude that STAT1/2 binding (representing bona fide ISGF3 complexes together with IRF9) at promoters was the main driver of ISG activation in our system (*n* = 71 in ESCs; *n* = 112 in MEFs). In addition, ISG activation was provided for a smaller fraction of promoters by STAT2 in the absence of STAT1, in line with the conclusion that the STAT2-IRF9 complex alone could provide some activation (Platanitis et al, 2019) (*n* = 5 in ESCs; *n* = 34 in MEFs). STAT1 without STAT2 appeared to lack significant activation capacity in our system but rather displayed some propensity to bind to already active promoters. Nevertheless, it could potentially be involved in promoting transcription of some ISGs where it was found at the promoter (*n* = 10 in ESCs; *n* = 11 in MEFs). For a remaining fraction of 105 (ESCs) and 306 (MEFs) ISGs, no STAT binding at the promoter was detected. Accordingly, these ISGs were either secondary target genes or become activated from non-promoter STAT binding sites. Based on these findings, we focused on STAT1/2 binding sites as a proxy for the ISGF3 complex to further characterize the relation between non-promoter STAT1/2 binding and ISGs.

### STAT1/2 enhancers are predicted from co-accessibility analysis

The non-promoter STAT1/2 peaks at intronic or intergenic sites could represent enhancer elements that regulate ISGs from a distance. A simple assignment of these potential enhancer sites to the nearest gene linked these sites to only a few additional ISGs that lacked promoter bound STAT1/2 (*n* = 13 in ESCs; *n* = 41 in MEFs) (**Supplementary Fig. 4A**). Thus, the assumption that the majority of enhancer targets can be predicted by selecting the closest gene appeared not be justified in our system. To further characterize potential targets of STAT1/2 binding at putative enhancers, we applied a novel strategy to define co-regulatory sites using co-accessibility events from single-cell ATAC (scATAC-seq) data (**Fig. 4A-C, Supplementary Fig. 4B, Supplementary Table S3**). The single-cell embeddings of chromatin accessibility showed no clear separation of ESCs and MEFs after IFNβ treatment (**Fig. 4A**), indicating that the observed gain in chromatin accessibility at STAT1/2 sites was not accompanied by a global alteration of the chromatin landscape (**Supplementary Fig. 4B**). These observations were in agreement with the RNA-seq data, where a couple of hundred ISGs were identified. For MEFs, two separate cell clusters were detected (**Fig. 4B**) and assigned to epithelial- and mesenchymal-like MEF subtypes by integration with the scRNA-seq data (**Fig. 1D, Fig. 4C**). Next, we computed correlations between pairs of genomic loci that were simultaneously accessible in the same cell based on previously described approaches (Granja et al, 2021; Mallm et al, 2019) to detect enhancers with STAT1/2 linked to ISGs before and after IFNβ stimulation (**Supplementary Fig. 4C, Supplementary Data Set 3**). This analysis was conducted for 392 STAT1/2 binding sites in ESCs and MEFs with all 2 kb peaks in a surrounding 1 Mb region.

**Figure 4.**
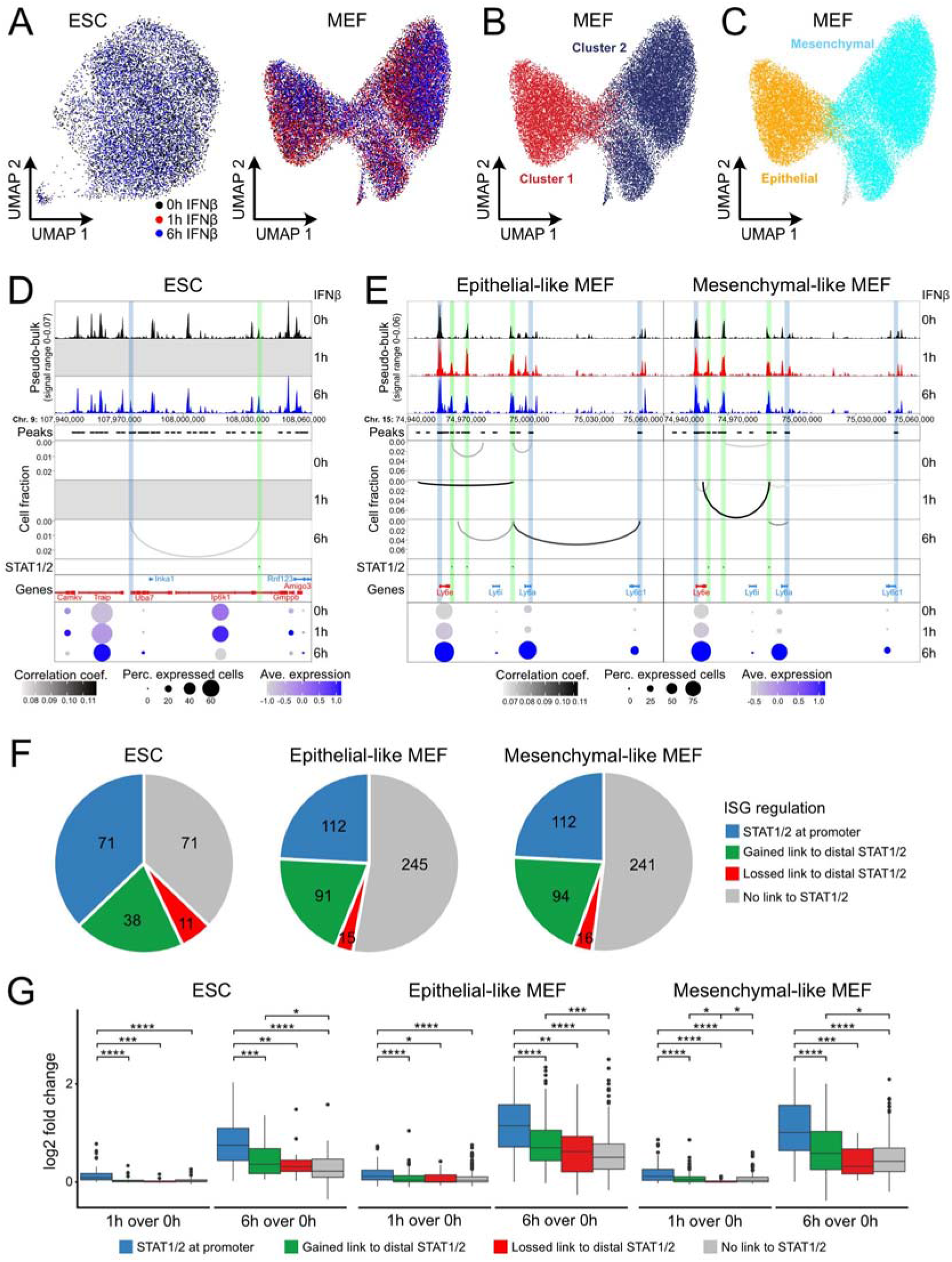
Regulation of ISG expression by distal STAT1/2 binding. (**A**) Single-cell embedding of chromatin accessibility in ESCs and MEFs with coloring according to treatment. (**B**) Same as panel *A* for MEFs with coloring according to clusters predicted by ArchR/Seurat. (**C**) Same as panel B with coloring according to MEF subtypes according to intregrated scRNA-seq data. (**D**) Co-accessibility maps before in ESCs of a region around the *Uba7* ISG. Top: Browser tracks of pseudo-bulk chromatin accessibility from single cells. Middle: Co-accessible links between the indicated intronic STAT1/2 site and other genomic loci. Bottom: Gene expression levels from scRNA-seq. Experimentally identified ISG promoters and STAT1/2 binding sites were marked by blue and green vertical bars, respectively. Transcription from the *Inka1* and the *Rnf123* gene was not detected. (**E**) Same as panel D but for three intergenic STAT1/2 binding sites in the *Ly6* ISG cluster in MEFs. (**F**) ISG regulation by STAT1/2 binding in ESCs (left), epithelial-like (mid), and mesenchymal-like MEFs (right). (**G**) Expression changes of ISGs for the different STAT1/2 dependent regulation types in ESCs (left), epithelial-like (mid), and mesenchymal-like MEFs (right).

As an exemplary result, induction of the *Uba7* ISG by STAT1/2 binding to a putative distal enhancer in ESCs is depicted in **Fig. 4D**. The IFNβ-induced co-accessible link between the STAT1/2 bound enhancer candidate and the *Uba7* promoter was associated with an increase in *Uba7* expression in scRNA-seq. *Uba7* was previously identified by bulk RNA-seq data as an ISG (**Fig. 1B**). Another example of ISG regulation by STAT1/2 binding to distal putative enhancers was the gene cluster of the Ly6 family in MEFs (**Fig. 4E**). In the gene cluster, expression of ISGs *Ly6e, Ly6a* and *Ly6c1* increased with IFNβ treatment. The promoter of ISG *Ly6e* was highly accessible before and after IFNβ treatment while *Ly6a* and *Ly6c1* promoters remained in relatively lowly accessible states. In contrast, the three intergenic STAT1/2 binding sites in this genomic region opened up strongly upon IFNβ treatment. Multiple co-accessible links between the intergenic STAT1/2 sites and ISGs were detected, either directly to the *Ly6* promoters or indirectly to their gene bodies or proximal regions. These involved the formation of new links between the potential enhancer cluster and the *Ly6a* and *Ly6c1* promoters as well as the loss of links present at the 0 h time point. With this co-accessibility analysis, we were able to link roughly 25% of ISGs without STAT1/2 promoter binding to a distal STAT1/2 binding event (**Fig. 4F**) (ESCs, 38 ISGs; epithelial-like MEFs, 91 ISGs; mesenchymal-like MEFs, 94 ISGs) (**Supplementary Data Set 3**). Interestingly, we also observed a loss of existing co-accessible links between ISGs and distal STAT1/2 sites at several loci (ESCs, 11 ISGs; epithelial-like MEFs, 15 ISGs; mesenchymal-like MEFs, 16 ISGs), which points to larger changes of the 3D chromatin organization during activation that could involve the resolution of inhibitory interactions.

### Binding of STAT1/2 to distal sites efficiently induces target ISG expression

Next, we investigated the expression induction for the differently regulated ISG categories after 1 h and 6 h of IFNβ treatment over unstimulated control cells and found similar patterns for ESCs and both MEF subtypes (**Fig. 4G**). After 1 h of IFNβ treatment some induction was observed for all cell types and ISG categories. ISGs with a STAT1/2 site at their promoter showed the strongest expression upregulation after 6 h of IFNβ treatment, which was significantly stronger than expression induction in all other ISG regulation categories. Additionally, ISGs that gained a co-accessible link to a distal STAT1/2 site showed a significantly stronger expression induction after 6 h of IFNβ treatment compared to ISGs without any link to STAT1/2. Moreover, the ISGs with a loss of a preexisting link to a distal STAT1/2 site upon IFNβ treatment showed a significantly lower gene expression level before IFNβ treatment (0 h) in ESCs and mesenchymal-like MEFs (**Supplementary Fig. 4D**). In summary, the scATAC-seq data allowed us to distinguish different mechanisms by which STAT1/2 induces ISG expression from distal sites. It suggests that STAT1/2 induced ISG expression from distal enhancers in addition to binding directly at their promoters. Moreover, our analysis suggests that the loss of pre-existing links during STAT1/2 binding could be associated with the removal of inhibitory interactions.

### Five different chromatin states of STAT1/2 binding sites can be distinguished

The overlap of STAT1/2 peaks from ESCs and MEFs revealed 92 shared binding sites mostly at promoters (70/92). The 116 ESC-specific and 184 MEF-specific sites were predominantly at nonpromoter loci (100/116 and 118/184) (**Supplementary Fig. 5A**). We reasoned that the cell type specific STAT1/2 binding was dependent on the chromatin context.

Accordingly, we mapped six histone modifications (H3K4me1, H3K4me3, H3K9ac, H3K27ac, H3K9me3 and H3K27me3) by ChIP-seq as well as chromatin accessibility by ATAC-seq. Exemplary regions for ESCs and MEFs were shown (**Fig. 5A**). STAT1/2 binding at the *Usp18* promoter induced the gene in both cell types from a transcriptionally repressed to an active state. In contrast, *Ifi27* induction was apparent only in ESCs as compared to a constitutively active state in MEFs while *Gbp6* became active in MEFs and remained silent in ESCs. Of note, several additional ISRE motifs did not display STAT1/2 binding illustrating the requirement for a permissive chromatin state (**Fig. 5A**). To reveal chromatin features that are linked to STAT1/2 binding, normalized read counts in a window of ±1 kb around the peak center were computed for the different readouts (**Supplementary Fig. 5A**). These data were then subjected to unsupervised k-means clustering (**Fig. 5B, Supplementary Fig. 5B, C**). Five main clusters emerged that were annotated based on the combination of enriched chromatin features (**Fig. 5B**): (i) “ Active Promoter” was enriched for H3K4me3, H3K9ac and H3K27ac (Ernst et al, 2011). (ii) “Active Enhancer” was marked by H3K4me1 and H3K27ac (Creyghton et al, 2010). (iii) The “Bivalent” state carried active marks like H3K4me3 and repressive marks like H3K27me3 at the same time (Bernstein et al, 2006). (iv) The “Poised” state showed only H3K4me1 (Creyghton et al, 2010). (v) “Repressed” was marked by enrichment of H3K9me3 or H3K27me3 (Lehnertz et al, 2003; Morey & Helin, 2010).

**Figure 5.**
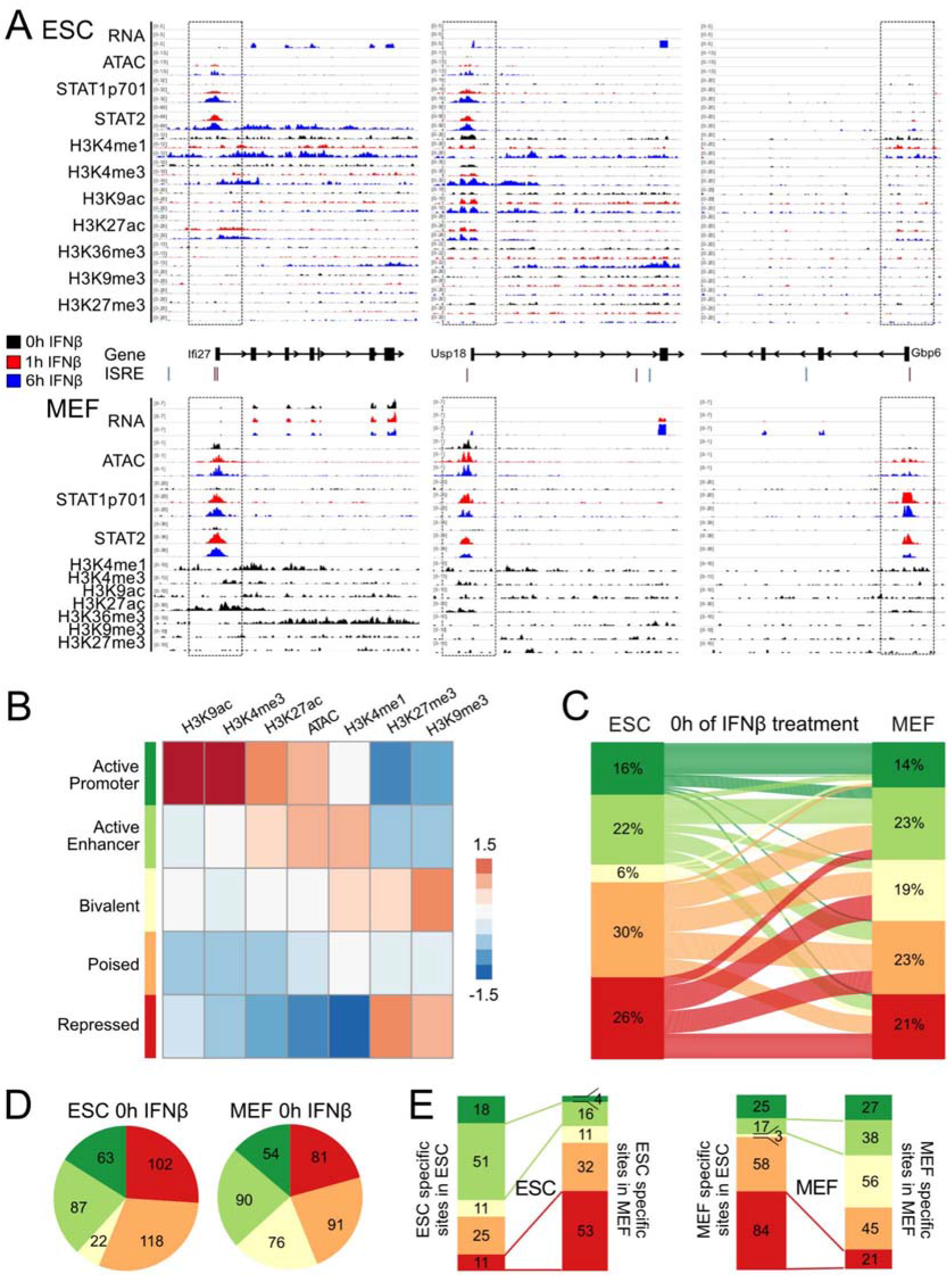
Contribution of chromatin features to STAT1/2 binding. (**A**) Genomic regions around the ISGs *Ifi27, Usp18* and *Gbp6* in ESCs (top) and MEFs (bottom) with the different sequencing readouts and the promoter regions marked by boxes. Gene annotation was based on ENSEMBL and the positions of the DNA binding motif IRSE were extracted from the HOMER database. Each browser track shows one representative biological replicate. (**B**) Heatmap of unsupervised k-means clustering of histone modifications and ATAC data at 392 STAT1/2 binding sites. The indicated five main chromatin states were identified. Data from unstimulated ESCs and MEFs as well as ESCs, at 1 h and 6 h IFNβ treatment were used. (**C**) Chromatin state transitions between untreated ESCs and MEFs at STAT1/2 binding sites based on the data in panel B and corresponding coloring of the five different chromatin states. (**D**) Absolute numbers of STAT1/2 binding sites according to chromatin states in unstimulated ESCs and MEFs. (**E**) Distribution of 116 ESC-specific and 184 MEF-specific STAT1/2 binding sites according to chromatin state.

### STAT1/2 binding is directed by chromatin accessibility and specific histone marks

Next chromatin states at STAT1/2 sites in ESCs and MEFs and their changes were analyzed (**Fig. 5C-E**). The most pronounced chromatin state transitions between cell types occurred from the “Poised” and “Repressed” states in ESCs to the “Active Enhancer”, “Bivalent” and “Poised” states in MEFs (**Fig. 5C**). The 116 ESC-specific sites displayed a 3 to 4-fold loss of the “Active Promoter” and “Active Enhancer” states and an approximately 5-fold increase of the “Repressed” state when their chromatin state was evaluated in MEFs (**Fig. 5E**). Corresponding changes of the “Active Enhancer” and “Repressed” states were also found for MEF-specific sites in ESCs and MEFs. The fraction of MEF-specific STAT1/2 sites in the “Active Promoter” state remained mostly unchanged between cell types, while the number of sites in the “Bivalent” state strongly increased from 3 to 56 sites (**Fig. 5E**). We conclude that the main changes that determine the cell type specific binding of STAT1/2 occurred between the “Repressed” state (H3K9me3 and H3K27me3) and “Active Enhancer” and “Bivalent” states that both are enriched in the H3K4me1 and H3K27ac modifications. Accordingly, the increased number of ISGs detected in MEFs appears to be related to the more frequent activation of ISRE containing enhancer elements.

To further dissect the relation between chromatin signals in the uninduced state and STAT1/2 binding upon induction we computed their correlations. Normalized read counts of a given chromatin feature before induction were plotted against STAT1/2 binding as represented by the average signal of STAT1 and STAT2 at 1 h of IFNβ treatment at the same locus (**Fig. 6A**). These plots visualized the differences between ESC-specific (black) and MEF-specific (red) binding sites for specific chromatin features. The *p*-value and correlation coefficient *R* of a given mark with STAT1/2 binding are plotted in **Fig. 6B**. ATAC (ESC, *R* = 0.42; MEF, *R* = 0.53), H3K4me1 (ESC, *R* = 0.45; MEF, *R* = 0.43) and H3K27ac (ESC, *R* = 0.41; MEF, *R* = 0.63) were the most strongly positively correlated marks, while H3K27me3 (ESC, *R* = −0.23; MEF, *R* = −0.39) was anticorrelated with STAT1/2 binding. For the repressive H3K9me3 mark the correlation was negative for ESCs (*R* = −0.26) and slightly positive for MEFs (*R* = 0.08) pointing to a more complex relation. We concluded that a pre-existing active chromatin state (open chromatin, H3K4me1 and H3K27ac) promoted STAT1/2 binding while chromatin loci marked by the H3K27me3 impeded this interaction (**Fig. 6C**).

**Figure 6.**
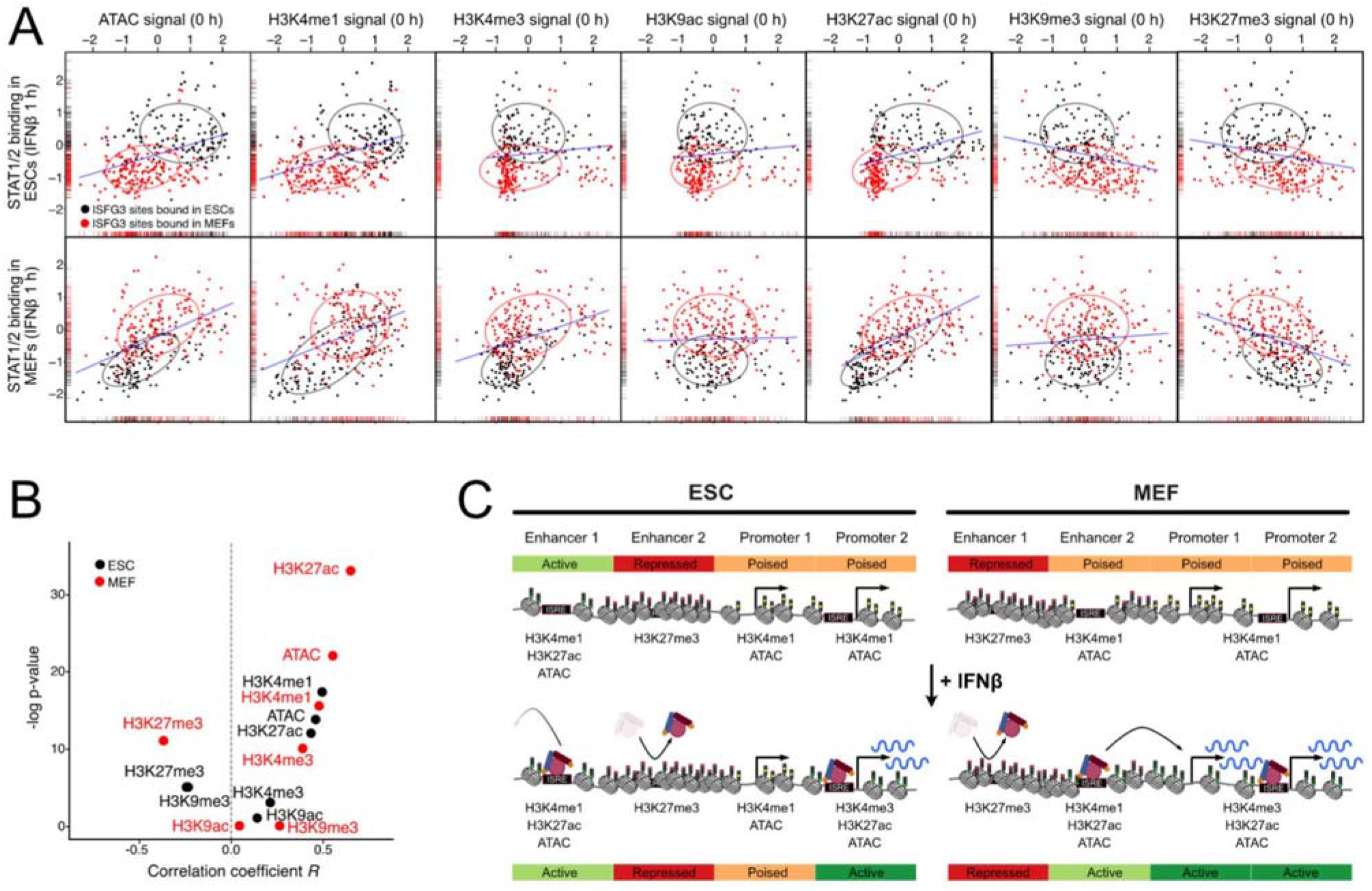
Correlation of STAT1/2 binding with pre-existing chromatin environment. (**A**) Correlation between STAT1/2 binding after 1 h of IFNβ treatment and pre-existing chromatin features before IFNβ treatment. The STAT1/2 binding signal was computed as the average signal of STAT1 and STAT2 after 1 h IFNβ treatment in ESCs (top) and MEFs (bottom). The chromatin features were quantified by counting the normalized read counts at the STAT1/2 binding sites before induction. ESC-specific STAT1/2 binding sites are shown in black and MEF-specific ones in red. Ellipses indicate the area, in which 75% of all data points are located. Density distributions are shown along the x- and y-axis. The blue line shows the linear regression of the combined set of ESC- and MEF-specific STAT1/2 binding sites. (**B**) Correlation between STAT1/2 binding and chromatin features determined for the data in panel A in ESCs (black) and MEFs (red). (**C**) Scheme of cell type specific ISG induction via STAT1/2 binding. The most prominent differences between cell types are depicted. Enhancers as well as promoters direct STAT1/2 binding in dependence of cell type specific chromatin states. Some ISGs have accessible ISRE motifs at their promoter and can directly be activated by STAT1/2 binding while a repressive chromatin state impedes binding at other promoters. The same applies to ISGs that lack an ISRE at the promoter but are activated by STAT1/2 binding to enhancers that induces transcription of a target ISG from a distal site.

## Discussion

We compared ESCs to *in vitro* differentiated NPCs as well as MEFs from the same mouse strain to reveal mechanisms that govern the cell type specific response to IFNβ in a genome wide manner. In total 513 ISGs were identified, in line with previous studies that reported between 200 to 1,000 upregulated genes in different cellular systems (de Veer et al, 2001; Der et al, 1998; Mostafavi et al, 2016). Our results corroborate the finding that ESCs show an attenuated response to IFNβ (D’Angelo et al, 2016; Gonzalez-Navajas et al, 2012; Guo, 2017; Guo et al, 2015; Wang et al, 2014; Wang et al, 2013; Whyatt et al, 1993). This stem cell specific feature appears to be compensated by a constitutive expression of some ISGs in human stem cells (Wu et al, 2018) as well as the presence of an antiviral RNA interference based system in mouse ESCs (Maillard et al, 2013; Poirier et al, 2021). A previous RT-qPCR analysis of selected components of the IFN signaling pathway in ESCs identified a significant downregulation of the IFNα/β receptor *Ifnar1* while *Stat2, Tyk2* and *Irf9* were upregulated as compared to a MEF cell line (Wang et al, 2014). Based on our differential RNA-seq maps of the unstimulated cell types we confirm the downregulation of *Ifnar1* in ESCs while the differences for *Stat2, Tyk2* and *Irf9* were above the p < 0.01 significance level. We additionally detected a strong downregulation of *Ifnar2*, the *Ifngr1/2* and the *Jak1/2* kinases in ESCs relative to NPCs and MEFs. Furthermore, both STAT1 and STAT2 as well as phosphorylated STAT1 were more abundant in MEFs than in ESCs at the protein level after induction. Thus, a globally reduced IFNβ response could be assigned to lower levels of key components of the IFN signaling pathway in ESCs.

Previous studies on STAT1/2 binding reported 6,703 STAT2 peaks for IFNα treated B cells (Mostafavi et al, 2016), and 41,582 (IFNγ-stimulated) and 11,004 (unstimulated) STAT1 binding sites in HeLa S3 cells (Robertson et al, 2007). The specificity of STAT peak assignment in these previous studies appears to be moderate. A fraction of 46% of the STAT2 peaks displayed a >2-fold increase upon IFNα treatment (Mostafavi et al, 2016), while a 2-5 fold enrichment of GAS and ISRE sequences in the STAT1 peaks was present (Robertson et al, 2007). Our identification of STAT1_p701_ and STAT2 binding sites was more stringent and displayed an at least 4-fold STAT enrichment upon induction. In addition, 80-90% of the sites carried a STAT- or IRF-family binding site sequence motif with a more than 10-fold higher frequency than found in the background sequences. It is further noted that we did not detect STAT2 ChIP-seq peaks before the IFNβ stimulus. Thus, an activity of unphosphorylated STAT2-IRF9 for basal gene expression of ISGs as reported in (Blaszczyk et al, 2015) was not apparent in the STAT2 binding maps recorded here.

The main ISG activation sites in our system had STAT1 and STAT2 bound simultaneously most likely within the ISGF3 complex that additionally involves IRF9 and in line with previous findings (Au-Yeung et al, 2013; Lee & Young, 2013; Singh et al, 2014). This assignment was confirmed by the binding motif analysis that yielded an enrichment of IRF motifs in 80-90% of the 392 STAT1/2 peaks. The number of sites that had only STAT1_p701_ or STAT2 bound was 1,037 and 323, respectively (**Supplementary Table S2**). STAT1 homodimers can also act as activators of type I IFN response (Stanifer et al, 2019; Stark & Darnell, 2012). However, the promoters that only had STAT1_p701_ bound showed no enrichment for ISGs in our data set. An additional minor contribution to ISG activation arose from binding of STAT2 in the absence of STAT1 at ISG promoters, which is in line with the observation that the STAT2-IRF9 complex has some activation capacity without STAT1 (Platanitis et al, 2019).

Interestingly, more than 2/3 of the STAT1/2 peaks were located at intergenic or intronic regions and thus represent potential enhancer elements that could drive ISG activation. The target ISGs of these putative enhancers did not appear to be those that were in closed genomic distance. We identified links between ISG promoters and distal STAT1/2 binding sites by applying a co-accessibility analysis of the scATAC-seq data (Granja et al, 2021; Mallm et al, 2019). This approach exploited the accessibility information obtained from thousands of single cells to compute pair-wise correlations of sites with bound STAT1/2 and ISG promoters. These correlations could originate from direct spatial contacts or other mechanisms of enhancer-promoter communication (Karr et al, 2022). With this approach we were able to link 25% of ISGs lacking STAT1/2 at the promoter to a distal STAT1/2 binding site that is likely to represent an enhancer that activates this gene. In addition, our data suggest that IFNβ induction and STAT1/2 binding could also involve the removal of pre-existing inhibitory links between ISGs and distal regulatory regions. The latter process might be related to the loss of long-range interactions observed during induction of differentiation in mouse ESCs (Feldmann et al, 2020). Furthermore, a recent study describes the reorganization of the 3D genome around ISG loci upon both IFNβ and IFNγ treatment, which involves loop formation, nucleosome remodeling and an increase of DNA accessibility (Platanitis et al, 2021). Thus, it is emerging that a reorganization of long-range chromatin interactions represents an important part of IFN mediated gene induction.

According to the HOMER data base (Heinz et al, 2010), ISREs are and are found at 134,069 loci in the mouse genome. According to our ChIP-seq analysis a much lower number of 392 ISREs had STAT1/2 bound. This large difference led us to characterize their chromatin environment as a determinant of STAT binding via a genome-wide correlation analysis. We find that a repressive chromatin conformation marked by H3K27me3 renders ISREs less accessible to STAT1/2 binding. In contrast, H3K4me1 and H3K27ac as well as an open chromatin state detected by ATAC were associated with sites permissive for STAT1/2 binding. These results are in line with a previous study that compared histone modifications at 18 ISREs (Testoni et al, 2011). In the latter data set, 6 out of 9 ISREs at activated promoters showed some enrichment for H3K4me1 before induction with IFNα. It is noted that H3K4me1 has been related to targeting the BAF (SWI/SNF) chromatin remodeler to chromatin, which interacts with STAT1_p701_ and STAT2 via its BRG1 component. Accordingly, this histone modification could promote chromatin opening and subsequent STAT1/2 binding (Christova et al, 2007; Huang et al, 2002; Local et al, 2018).

## Conclusions

Our integrated multi-omics data set provides insight into the interplay between the IFNβ mediated activation of ISGs, STAT binding and chromatin features. It revealed a number of links that could be exploited to modulate the IFN response during virus infection or therapeutic intervention in cancer (Borden, 2019; Hoffmann et al, 2015). Numerous so called “epigenetic drugs” that inhibit enzymes setting or removing histone acetylation and methylation are already used in anti-cancer therapy (Cheng et al, 2019; Mohammad et al, 2019). In the light of our study, the resulting perturbances of chromatin features are also likely to affect IFN response. For example, histone deacetylase (HDAC) inhibitors that result in the hyper-acetylation of histones and render chromatin more accessible could enhance STAT1/2 binding to otherwise occluded ISREs (Cusack et al, 2020; Gorisch et al, 2005; Shogren-Knaak et al, 2006; Wang & Hayes, 2008). At the same time, however, these drugs also affect the acetylation state of protein factors involved in IFN mediated signaling like the acetylation and activity of the STAT1/2 complex itself (Tang et al, 2007). Accordingly, HDACs have been shown to both repress or enhance IFN response in a complex manner (Au-Yeung & Horvath, 2018; Lu et al, 2019). Thus, changing STAT1/2 binding patterns more specifically would require a targeted approach beyond global inhibition/activation of epigenetic modifiers like HDACs. This could be achieved, for example, by using more selective drugs (Cheng et al, 2019; Mohammad et al, 2019) or dCas9 mediated changes of ISRE chromatin states at promoters and enhancers by targeted binding of activators that sets or remove H3K27ac, H3K4me1 or H3K27me3 (Erdel et al, 2020; Frank et al, 2021; Li et al, 2020). In this manner, ISG activation patterns could be changed to modulate the cell type specific antiviral response.

## Materials and methods

### Cell culture work and IFNβ treatment

Mouse 129/Ola ESCs, NPCs differentiated *in vitro* from ESCs and MEFs were cultured at 37 °C with 5 % CO_2_ and routinely checked for the absence of mycoplasma contaminations as described previously (Bibel et al, 2007; Mallm et al, 2020; Teif et al, 2012). IFNβ was prepared from a BHK cell line over-expressing IFNβ and grown with DMEM supplemented with 10 % FCS, 1 % L-glutamine and 1% penicillin/streptomycin. After growing the cells in the same medium but with 2% FCS for 24 h the IFNβ containing medium was passed through a 0.45 μm sterile filter and stored in aliquots at −80°C. The resulting IFNβ stock was calibrated against a commercial IFNβ preparation (Sigma) by using a Mx2-luciferase reporter cell line (Schwerk et al, 2013). A stock concentration of 16.6 U/μl was calculated. For induction, cells were treated with IFNβ at a concentration of 500 U/ml for 1 hour or 6 hours.

### Western blots

Western blot samples were prepared by collecting cells directly out of cell culture. Cells were transferred into 1.5 ml tubes, washed once with PBS and counted. A 50 μl volume of pre-prepared RIPA buffer (150 mM NaCl, 1 % NP40, 50 mM Tris-HCl, pH 8.0, 0.5 % sodium deoxycholate, 0.1 % sodium dodecyl sulfate) were added per 0.5 million cells in suspension. The mixes were incubated for 60 min on ice, spun down at max speed at 4 °C for 30 min. Supernatant was transferred to a fresh tube and stored at −20 °C. Gels were blotted on LF PVDF membranes using the trans-blot turbo transfer system (Bio-Rad) and blocked with 5 % BSA in Tris-buffered saline supplemented with 0.1% Tween 20 detergent (TBST) at room temperature for 1 h. The primary antibodies were diluted according to manufacturer’s recommendations in 5 % BSA and incubated at 4 °C. On the following day, the membrane was washed three times with TBST buffer at room temperature for 5 min under agitation and incubated with secondary anti-HRP antibody (normally 1:5000 diluted in 5 % BSA) at room temperature for 1 h, washed three times with TBST, incubated with clarity western ECL substrate for 5 min and imaged. The antibodies used are listed in **Supplementary Table S4**.

### Bulk RNA-seq

Cells were seeded on a 6-well plate. Two (ESCs and MEFs) or five (NPCs) days after plating, cells were washed two times with PBS. Then 500 μl LBP was added and cells were stored at −80 °C. RNA was isolated with the Nucleospin RNA kit (Macherey-Nagel) according to the manufacturer’s protocol. The elution step was done two times with 30 μl RNase-free water within the same tube.

Concentrations were measured by Qubit RNA HS assay kit and the quality of RNA was analyzed on Tapestation D5000 HS (Agilent). Removal of rRNAs from isolated samples of IFNβ stimulated ESCs and MEFs was done following the protocol of Ribo-Zero rRNA removal kit. An input of 5 μg total RNA was used and the depleted RNAs were eluted in 30 μl Rnase-free water supplemented with 1 μl RiboLock Rnase-inhibitor (40 U/μl). Concentrations were measured by Qubit RNA HS assay kit. For NPCs, RNA samples were treated with Dnase at 37 °C for 30 min and purified by ethanol precipitation. Concentrations were measured by Qubit RNA HS assay kit and 750 ng of Dnase-treated RNA was used for rRNA depletion by NEB Next rRNA depletion kit (Human/Mouse/Rat). The depletion was performed based on the manufacturer’s protocol. Samples were purified with RNA Clean XP beads (Beckman) with a 2.2x ratio and finally eluted in 8 μl nuclease-free water. Purified rRNA-depleted RNA samples of ESCs, MEFs and NPCs were used to prepare NGS libraries based on the NEB Next Ultra II directional RNA library preparation kit from Illumina. As default, 50 ng of rRNA-depleted RNA was used as input. For less concentrated samples, 10 ng were used. The RIN value of all samples were above 7 and therefore the mixes were incubated at 94 °C for 15 min. Further, a 5-fold NEB Next adaptor dilution was used as default at the adaptor ligation step. For lower concentrated samples, a 25-fold dilution was used. All samples were dual-barcoded with unique i5 and i7 primers. For 50 ng samples a total of nine cycles and for 10 ng samples eleven cycles were performed during the PCR enrichment of the adaptor ligation DNA step. Samples were measured with Qubit dsDNA HS assay kit and the fragment size was determined with a Tapestation D5000. In total, four replicates of ESCs, two replicates of MEFs and four replicates of NPCs treated with IFNβ for 0 h, 1 h and 6 h were acquired. The corresponding RNA-seq libraries were 50-bp single-end sequenced on the HiSeq 4000 System (Illumina) with at least 50 million reads per sample. Sequencing of RNA, as well as that of all other sequencing readouts, was done at the DKFZ Genomics and Proteomics Core Facility.

### ChIP-seq of STAT1_p701_ and STAT2

STAT1_p701_ and STAT2 ChIPs were performed with the ChIP enzymatic chromatin IP kit from Cell Signaling Technology according to the manufacturer’s protocol. Around 4 x 10^6^ cells per sample were used as input for the ChIPs. The Mnase digestion step was not used for ESCs and MEFs after formaldehyde fixation. Chromatin fragmentation was done with the Epishear probe sonicator (Active Motif) at 4 °C with 30 s long on and off cycles and 50 % amplitude. For ESCs and MEFs ten cycles of chromatin fragmentation were performed to yield an average fragment size of around 150 bp. The immunoprecipitation was conducted with 10 μg of chromatin in a total volume of 500 μl and addition of antibodies (**Supplementary Table S3**). The sequencing libraries were prepared using the NEB Next Ultra II DNA library preparation kit for Illumina with 40 μl ChIP sample and added 10 μl 10 mM Tris-HCl pH 8.0. For the input reaction, a 1:10 dilution was made and from this dilution 4 μg chromatin were used and filled up with 1x 10 mM Tris-HCl pH 8.0 to a total volume of 50 μl. Concentrations were measured by Qubit dsDNA HS assay kit and fragment distribution was analyzed on a Tapestation D5000. The libraries were sequenced as described above for RNA-seq.

### ChIP-seq of histone modifications

ESCs were cultured in 150 mm dishes and treated with IFNβ for 0 h, 1 h or 6 h. Media was removed and cells were detached with Accutase, washed with PBS supplemented with PMSF at 0.5 mM concentration and crosslinked with 1 % formaldehyde (1 ml 16 % formaldehyde with 15 ml PBS) for 10 min at room temperature. 125 mM glycine was added to neutralize the formaldehyde and rotated at room temperature for 5 min. Afterwards, samples were washed three times with PBS/100 mM PMSF and cell pellets were resuspended in 10 ml swelling Buffer (25 mM Hepes pH 7.8, 1 mM MgCl_2_, 10 mM KCl, 0.1 % NP-40, 1 mM DTT, 0.5 mM PMSF). A 10 min incubation step on ice and a centrifugation step at 2000 rpm for 5 min at 4 °C was performed. 4 x 10^7^ cells were resuspended in 100 μl Mnase Buffer and 40 U Mnase was added. After an incubation step at 37 °C for 15 min, 100 μl of 10x sonication buffer and 800 μl water were added. Samples were incubated on ice for 5 min, transferred into 12×24 mm tubes and sonicated for 15 min (burst 200; cycle 20 %; intensity 8) on a Covaris sonicator. A centrifugation step was performed at 13,000 rpm and 4 °C for 15 min. The supernatant was transferred into fresh tubes and chromatin was snap frozen with liquid nitrogen and store at −80 °C. A quality check of reverse cross-linked samples was performed and yielded a fragment size of around 150 bp for the sheared chromatin. Pre-equilibrated 25 μl protein G beads were used per sample at room temperature for 10 min in sonication buffer (10 mM Tris pH 8.0, 200 mM NaCl, 1 mM EDTA, 0.1 % Na-deoxycholate, 0.5 % n-lauroylsarcosine, 0.5 mM PMSF). A sample precleaning step was performed by adding 25 μl Protein G beads with 4 μg IgG antibody (rabbit or mouse) to chromatin and incubated rotating at 4 °C for 2 h. Beads were pelleted, and supernatant was transferred to fresh tubes. Antibodies were added to chromatin samples and incubated at 4 °C for 2 h. Then, 25 μl of pre-equilibrated beads were added to the samples and incubated rotating at 4 °C O/N. The beads were washed by rotating at 4 °C for 5 min with high-salt buffer (50 mM Hepes, pH 7.9, 500 mM NaCl, 1 mM EDTA, 1 % Triton-X-100, 0.1 % Na-deoxycholate, 0.1 % SDS, 0.5 mM PMSF), lithium buffer (20 mM Tris pH 8.0, 1 mM EDTA, 250 mM LiCl, 0.5 % NP-40, 0.5 % Na-deoxycholate; 0.5 mM PMSF) and 2x with TE-buffer (10 mM Tris pH 8.0, 1 mM EDTA). Each sample was eluted two times with 250 μl elution buffer (50 mM Tris pH 8.0, 1 mM EDTA, 1 % SDS, 50 mM NaHCO_2_) at 37 °C for 15 min on a shaker. Reverse cross-linking was performed by adding 20 μl 5 M NaCl and incubation at 65 °C overnight. 10 μl EDTA (0.5 M), 0.5 μl Rnase A (10 mg/ml) and 50 μl Tris (1 M, pH 6.8) was added and incubated at 37 °C for 30 min. Then, 2 μl Proteinase K (20 mg/ml) was added and incubated at 55 °C for 2 h. An isopropanol precipitation was performed to purify the DNA. Samples were resuspended in water. Samples were measured with Qubit dsDNA HS assay kit and the fragment size was determined on a Tapestation D5000. Libraries were sequenced as described above for RNA-seq. In ESCs, two replicates for H3K4me1, H3K4me3 and H3K27ac and three replicates for H3K9ac, H3K9me3 and H3K27me3 were sequenced. In MEFs, two replicates of all modifications were sequenced.

### Bulk ATAC-seq

ESCs were plated on 6 well plates and treated for 0 h, 1 h or 6 h with IFNβ at 500 U/ml. Cells were detached using accutase, collected and washed with 1xMT-PBS. A total of 50,000 cells were transferred into fresh tubes and centrifuged by 800 g at 4 °C for 5 min. For ESCs, the cell pellet was resuspended in 200 μl ATAC lysis buffer (10 mM Tris-HCl pH 7.4, 10 mM NaCl_2_, 3 mM MgCl_2_, 0.1 % NP-40), incubated at room temperature for 2 min and centrifuged at 800 g and 4 °C for 5 min. Supernatant was discarded and pellets resuspended in 20 μl ATAC reaction buffer containing 10 μl 2x transposase buffer and 2.5 μl Tn5 enzyme (Illumina). Samples were incubated at 37 °C for 30 min. Reactions were stopped by adding 5 μl EDTA (100 mM) in Tris-HCl pH 8.0 to final concentration of 20 mM. For MEFs and NPCs, the cells were directly resuspended in 25 μl ATAC reaction buffer with digitonin (9.75 μl H_2_O, 12.5 μl 2x transposase buffer (Illumina), 0.5 μl 50x proteinase inhibitor, 2 μl Tn5 enzyme (Illumina), 0.25 μl 1 % digitonin) and incubated at 37 °C for 30 min. The samples were purified with a MinElute PCR purification kit (Qiagen) and eluted in 12 μl buffer. After PCR amplifications, sequencing libraries were purified with AMPure beads (Beckman). Concentration was measured with the Qubit dsDNA HS assay kit (ThermoFisher) on a Qubit fluorometer, and size distribution of final library was checked on a Tapestation D5000. Libraries were 50-bp paired-end sequenced on Illumina HiSeq 2000 or 4000 systems with at least 50 million reads per sample. Two replicates for ESCs and NPCs and four for MEFs were sequenced.

### Analysis of bulk sequencing data

For RNA-seq analysis, ribosomal RNAs were removed and raw reads were mapped with STAR (Dobin et al, 2013) to the mm10 mouse reference genome and normalized read counts (transcripts per kilobase million, TPMs) were computed with RSEM (Li & Dewey, 2011). The differential gene expression analysis between treated and untreated controls was performed using DESeq2 (Love et al, 2014) with p-value <0.05 and log fold change >1.5. For the analysis of ChIP-seq and ATAC-seq data, reads were mapped with Bowtie2 (Langmead & Salzberg, 2012) to the mm10 mouse reference genome. Duplicates and reads annotated to blacklisted regions (Encode Project Consortium, 2012) as well as mitochondrial reads were removed. Quality control followed the Encode guidelines (https://www.encodeproject.org/data-standards/chip-seq/) and involved the fraction of reads in peaks (FriP) scores, normalized strand coefficients (NSC) and relative strand correlation (RSC) values for each sample. Peak calling was done with MACS2 (Zhang et al, 2008) and for H3K9me3 and H3K27me3 also with SICER (Xu et al, 2014) with p-value threshold of 10^-5^. STAT1_p701_ and STAT2 binding sites were identified against the unstimulated controls for 1 h and 6 h of IFNβ treatment from the ChIP-seq data with Diffbind (Ross-Innes et al, 2012) using the consensus peak list and thresholds of FDR <0.05 and 4-fold enrichment. Sequence motifs enriched in STAT1, STAT2 and STAT1/2 peaks were identified using HOMER (Heinz et al, 2010). For the analysis of the STAT1/2 chromatin environment, STAT1/2 bound sites in ESCs and MEFs were expanded by 1 kb up- and downstream. The ChIP- and ATAC-seq signal in these regions was determined from the respective read counts after normalizing for library depth and fragment length and computing enrichments over histone H3 for histone modifications and IgG for STAT1/2. Replicates of the same samples and time points of IFNβ stimulation were averaged. The resulting count tables were used as input for the k-means clustering to characterize the chromatin environment at STAT1/2 binding sites.

### Single-cell RNA-seq and ATAC-seq

The scRNA-seq experiments were performed based on the standard protocol for the Chromium single-cell 3’ reagent kit v2 (10x Genomics). ESCs and MEFs were treated for 0 h, 1 h or 6 h with IFNβ. The cDNA amplification was done by running 13 PCR cycles. The samples were eluted again in 35 μl 10 mM Tris-HCl pH 8.0. Concentrations of cDNA libraries were measured by Qubit dsDNA HS assay kit and mean peak sizes of the samples were determined on a Tapestation D5000. Each of the final libraries were paired-end sequenced (26 bp and 74 bp) on one Illumina HiSeq 4000 lane. For scATAC-seq, ESCs and MEFs were treated with IFNβ for 0 h, 1 h (only for MEFs) or 6 h and libraries were prepared according to the Chromium single-cell ATAC v1.0 protocol (10x Genomics). Two (three for MEFs treated with IFNβ for 6 h) lane replicates per scATAC libraries were paired-end sequenced on an Illumina NovaSeq 6000 system according to the manufacturer’s protocol.

### Analysis of scRNA- and scATAC-seq data

Sample demultiplexing and barcode processing of scRNA-seq data was conducted with the Cell Ranger pipeline from 10x Genomics. For ESCs, quality filtering was conducted by selecting only cells within a certain percentage of mitochondrial reads (2.5 % < accepted cells < 7.5 %) and number of detected genes (2,000 < accepted cells < 6,500), yielding 1,332 cells for time point 0 h, 2,085 cells for 1 h and 4,825 for 6 h of IFNβ stimulation. For MEFs, quality filtering was conducted by selecting only cells within a certain percentage of mitochondrial reads (0.5 % < accepted cells < 7.5 %) and number of detected genes (1,250 < accepted cells < 6,500), yielding 9,771 cells for time point 0 h, 10,186 cells for 1 h and 7,579 for 6 h of IFNβ stimulation. Further analysis was done using the R package Seurat (Stuart et al, 2019). The scATAC-seq data were demultiplexed and aligned with Cell Ranger ATAC count (10x Genomics) using the provided mouse mm10 reference. Further processing of the data was conducted with ArchR (Granja et al, 2021). Cells were filtered using a minimal and maximal threshold for number of fragments (10^3.5^ and 10^5^, respectively), a TSS ratio above 4 and a ratio of fragments in blacklisted genomic regions to all fragments below 0.0225 (ESCs) and 0.016 (MEFs). Co-accessibility between genomic regions was separately calculated for cell types and treatment conditions adjusting the ArchR framework to single-cell resolution without aggregation of cells. The degree of co-accessibility in the background was determined by randomly shuffling the accessibility values over cells and peaks as described previously (Mallm et al, 2019). The 99^th^ percentile of the maximum shuffled background co-accessibility score was used as a threshold to determine true co-accessible links. Co-accessible links were further evaluated by percent of accessible cells in the linked peak pairs.

## Data access

The data and computer code produced in this study are available from the following sources: All original sequencing and relevant processed data have been deposited under GSE160764 at Gene Expression Omnibus (https://www.ncbi.nlm.nih.gov/geo/). Software used for data analysis for the different sequencing readouts is listed in **Supplementary Table S5**.

## Author contributions

Study design: KR, MM, FE

Acquisition of data: MM, CB, KMO, JPM, JH

Analysis of data: MM, IL, KMO, FE, LK, NK

Drafting of manuscript: MM, KR

Manuscript reviewing: all authors

Supervision and coordination: KR

## Acknowledgments

We thank Jorge Trojanowski, Sabrina Schumacher and Katharina Bauer for discussions and help and Mario Köster and Hansjörg Hauser for providing the IFNβ over-expression cell line. This work was supported by grants TRR179 (Z03) and Ri1283/14-1 of the German Research Foundation (DFG) to KR. We thank the DKFZ High Throughput Sequencing and the Omics IT and Data Management core facilities for support and services and the Central Animal Laboratory for preparation of mouse fibroblast cells. Additional data storage at SDS@hd was supported by the Ministry of Science, Research and the Arts Baden-Württemberg and the DFG through grants INST 35/1314-1 FUGG and INST 35/1503-1 FUGG.

