## Supplementary Information for "Epigenetic signals that direct cell type specific interferon beta response in mouse cells"

### Content

#### Supplementary Figures

Figure S1. GO terms and differential gene expression analysis of nascent RNA and single-cell data.

Figure S2. Gene expression thresholds and expression of IFN signaling genes.

Figure S3. Cell type specific binding of STAT1 and STAT2.

Figure S4. Identification of co-accessible STAT1/2 binding sites and target genes.

Figure S5. Clustering of chromatin features at STAT1/2 binding sites.

#### Supplementary Tables

Table S1. Number and ID of biological replicates for different sequencing readouts.

Table S2. Cell type specific ISGs and STAT1/2 binding sites.

Table S3. Antibodies used in this study.

Table S4. Data analysis software.

Table S5. scATAC-seq data overview and quality.

Table S6. Inventory of supplementary data sets.

#### Supplementary Data Sets

Additional data sets on samples and the analysis results derived from the different sequencing readouts are provided as separate files in Microsoft Excel format. An inventory for these data sets is given in Supplementary Table S6.

#### Supplementary References

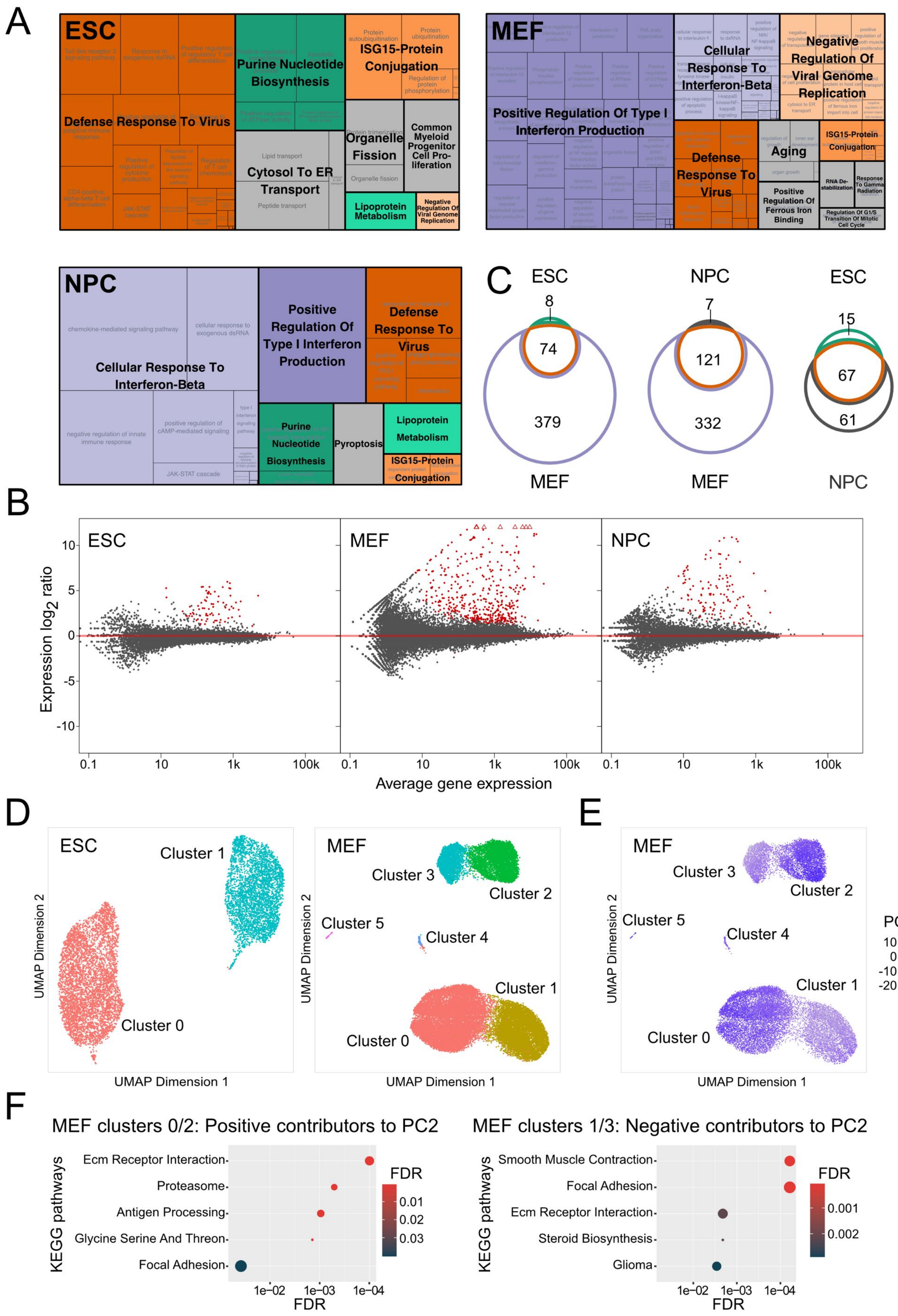
A
ESC
Defense Response To Virus
Purine Nucleotide Biosynthesis
ISG15-Protein Conjugation
Organelle Fission
Common Myeloid Progenitor Cell Proliferation
Cytosol To ER Transport
Lipoprotein Metabolism
Negative Regulation Of Viral Genome Replication
MEF
Cellular Response To Interferon-Beta
Negative Regulation Of Viral Genome Replication
Positive Regulation Of Type I Interferon Production
Defense Response To Virus
Aging
ISG15-Protein Conjugation
Positive Regulation Of Ferrous Iron Binding
RNA De-stabilization
Response To Gamma Radiation
Regulation Of G1/S Transition Of Mitotic Cell Cycle
NPC
Cellular Response To Interferon-Beta
Positive Regulation Of Type I Interferon Production
Defense Response To Virus
Purine Nucleotide Biosynthesis
Pyroptosis
Lipoprotein Metabolism
ISG15-Protein Conjugation
C
ESC
8
74
379
MEF
NPC
7
121
332
MEF
ESC
15
67
61
NPC
B
ESC
MEF
NPC
Expression log2 ratio
Average gene expression
D
ESC
Cluster 1
Cluster 0
MEF
Cluster 3
Cluster 2
Cluster 5
Cluster 4
Cluster 1
Cluster 0
UMAP Dimension 1
UMAP Dimension 2
E
MEF
Cluster 3
Cluster 2
Cluster 5
Cluster 4
Cluster 1
Cluster 0
PC2
UMAP Dimension 1
UMAP Dimension 2
F
MEF clusters 0/2: Positive contributors to PC2
KEGG pathways
Ecm Receptor Interaction
Proteasome
Antigen Processing
Glycine Serine And Threon
Focal Adhesion
FDR
MEF clusters 1/3: Negative contributors to PC2
Smooth Muscle Contraction
Focal Adhesion
Ecm Receptor Interaction
Steroid Biosynthesis
Glioma
FDR

**Supplementary Figure 1. GO terms and differential gene expression analysis of nascent RNA and single-cell data.**

(**A**) Enriched GO terms of all ISGs in ESCs, MEFs and NPCs. Functional annotations of differentially expressed gene lists were generated by DAVID (Huang da et al, 2009) to identify gene ontology (GO) terms. The visualization resulting terms was conducted with REVIGO (Supek et al, 2011). Orange colored terms were found in all three cell types, green terms in ESCs and NPCs and violet terms were identified in the differentiated cell types MEFs and NPCs. Terms in grey color were specific to a single cell type. (**B**) Analysis of intronic reads to identify differentially expressed genes on the nascent RNA level after 6 h of IFNβ treatment. Intronic reads were counted using HTSeq (Anders et al, 2015) and a modified GTF file containing only intronic sites was used for the differential gene expression analysis with DEseq2 (Love et al, 2014). Red dots represented differentially expressed genes at the level of $p_{adj} < 0.05$ and fold change ≥ 1.5. A total of 82 (ESCs), 128 (NPCs) and 453 (MEFs) genes were upregulated while no downregulated genes were detected. (**C**) Overlap of all (0 h vs 6 h and 0 h vs 1 h) ISGs detected by analysis of intronic reads detected in at least one differential gene expression analysis in ESCs, NPCs and MEFs. (**D**) Single-cell embedding of gene expression in ESCs (left) and MEFs (right). Coloring depicts cell clusters predicted by Seurat. (**E**) Single-cell embedding of gene expression in MEFs. Coloring depicts the score of principal component 2 (PC2) per cell. The plot shows that the clustering of MEFs into two groups was mainly driven by PC2. (**F**) Overrepresented KEGG pathways in positive contributors to PC2 (right, contribution > 0.025) and negative contributors to PC2 (left; contribution < -0.025).

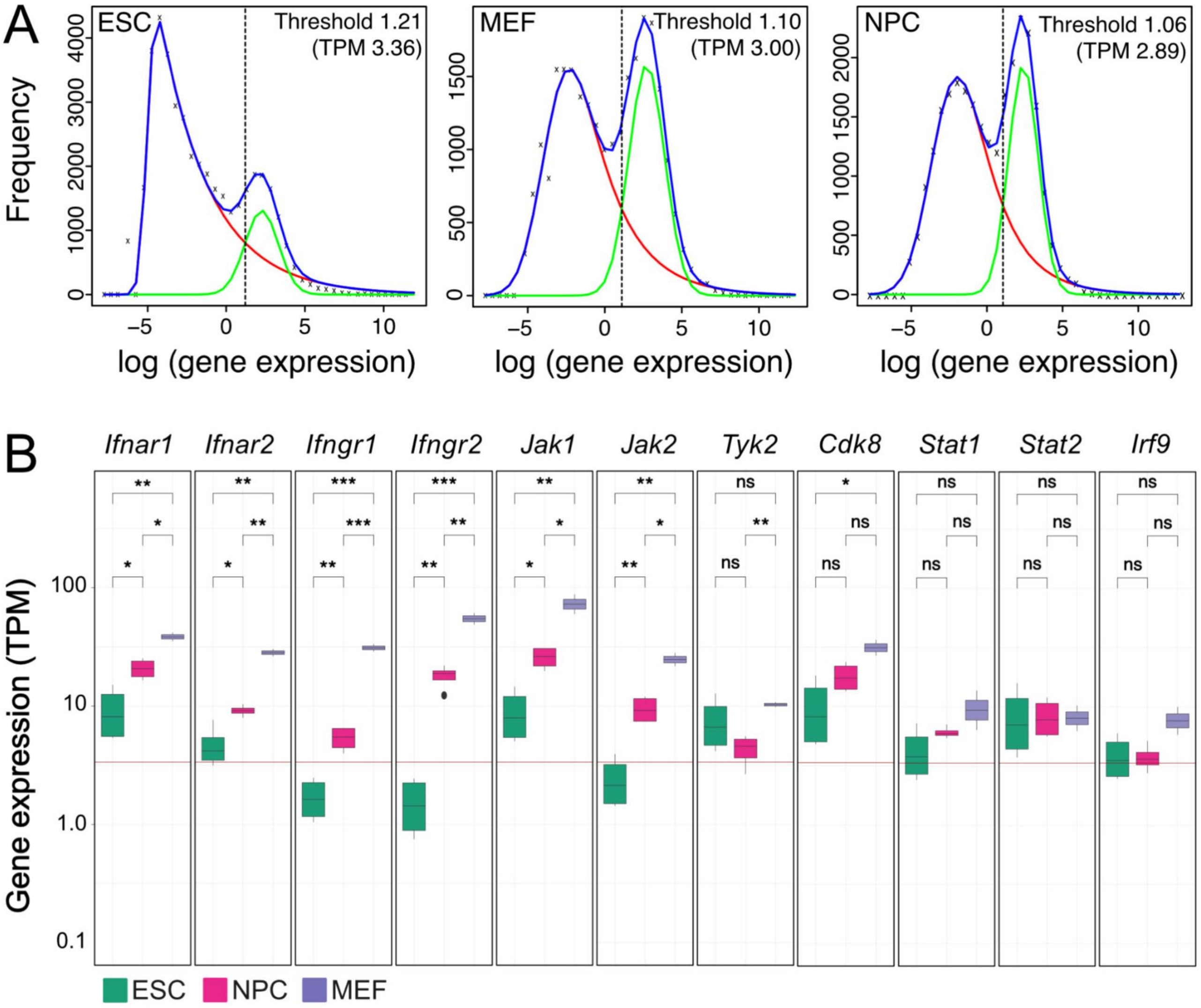


**Supplementary Figure 2. Gene expression thresholds and expression of IFN signaling genes.** (**A**) Normalized gene expression levels (log) at 0 h IFNβ in ESCs (left), MEFs (middle) and NPCs (right). Density curves of gene expression were shown in blue. Two Gaussian distributions were fitted to represent actively expressed (green) and repressed (red) genes. Their intersection points were marked by a dotted line and define the thresholds to distinguish actively expressed and repressed genes. (**B**) Normalized gene expression levels (TPM) of factors involved in IFN signaling. Gene expression levels of interferon receptors (*Ifnar1*, *Ifnar2*, *Ifngr1*, *Ifngr2*), JAK-STAT cascade kinases (*Jak1*, *Jak2*, *Tyk2*, *Cdk8*) and associated transcription factors (*Stat1*, *Stat2, Irf9*) were shown for unstimulated (0 h) ESCs (green), MEFs (purple) and NPCs (magenta). Significance was assessed with a paired t-test at the $p_{adj} < 0.05$ level. The red line represents the calculated threshold in ESCs to distinguish actively expressed and repressed genes.

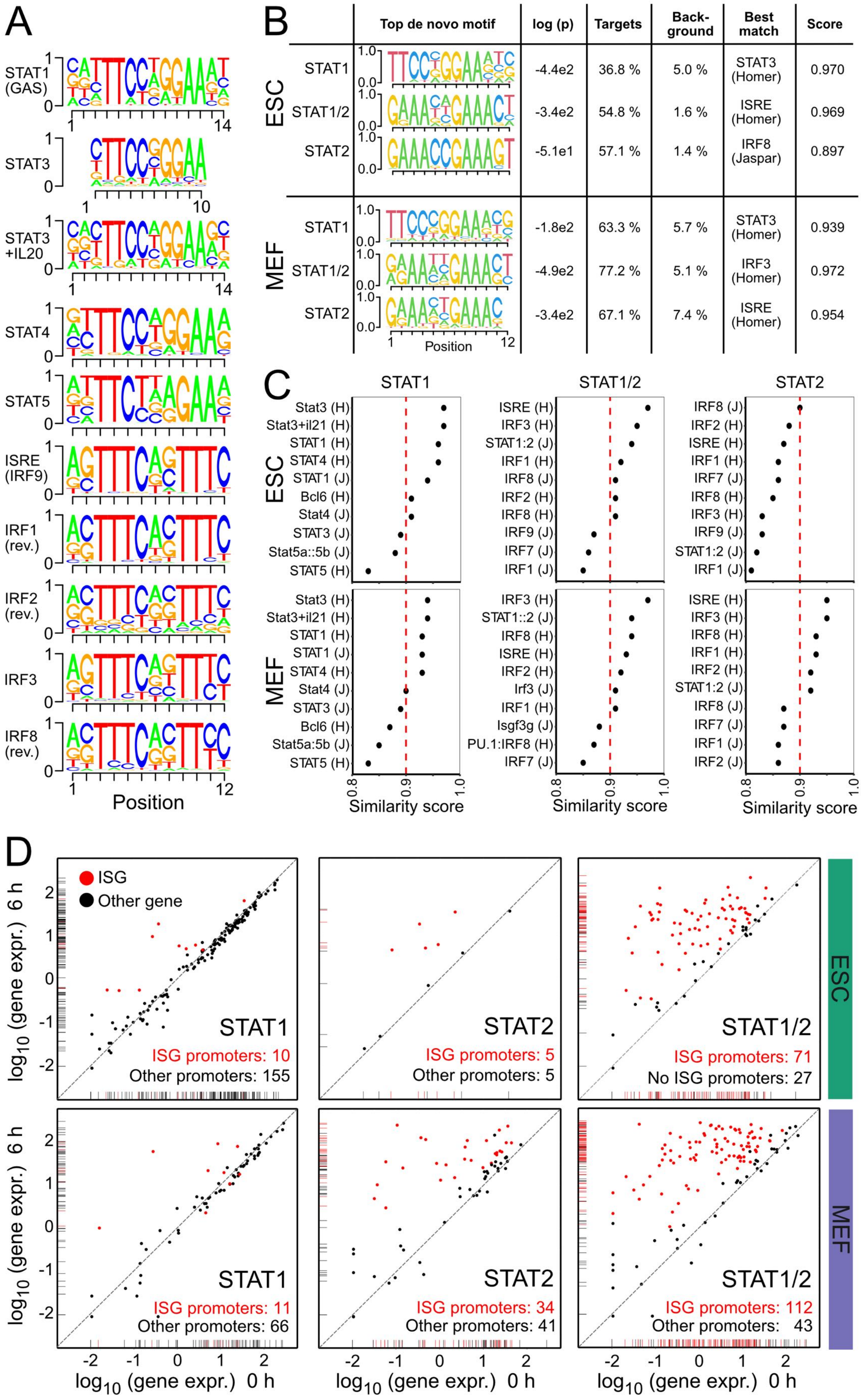
A
STAT1 (GAS)
STAT3
STAT3 +IL20
STAT4
STAT5
ISRE (IRF9)
IRF1 (rev.)
IRF2 (rev.)
IRF3
IRF8 (rev.)
Position
B
Top de novo motif
log (p)
Targets
Back-ground
Best match
Score
ESC
STAT1 -4.4e2 36.8 % 5.0 % STAT3 (Homer) 0.970
STAT1/2 -3.4e2 54.8 % 1.6 % ISRE (Homer) 0.969
STAT2 -5.1e1 57.1 % 1.4 % IRF8 (Jaspar) 0.897
MEF
STAT1 -1.8e2 63.3 % 5.7 % STAT3 (Homer) 0.939
STAT1/2 -4.9e2 77.2 % 5.1 % IRF3 (Homer) 0.972
STAT2 -3.4e2 67.1 % 7.4 % ISRE (Homer) 0.954
Position
C
STAT1
STAT1/2
STAT2
ESC
MEF
Similarity score
D
ISG
Other gene
STAT1
ISG promoters: 10
Other promoters: 155
STAT2
ISG promoters: 5
Other promoters: 5
STAT1/2
ISG promoters: 71
No ISG promoters: 27
ESC
STAT1
ISG promoters: 11
Other promoters: 66
STAT2
ISG promoters: 34
Other promoters: 41
STAT1/2
ISG promoters: 112
Other promoters: 43
MEF
log10 (gene expr.) 6 h
log10 (gene expr.) 0 h

**Supplementary Figure 3. Cell type specific binding of STAT1 and STAT2.**

(**A**) Position weight matrices (PWM) of top five STAT- and IRF-family motifs identified in $STAT1_{p701}$ and STAT2 ChIP-seq peaks based on HOMER annotation (Known motifs). PWM showed the probability of each nucleotide on the y-axis and the position within the motif on the x-axis. For IRF1, IRF2 and IRF8 the reverse complementary sequences were shown to be comparable with ISRE and IRF9 annotated motif. The source of the motifs was indicated with a letter behind the motif representing either the HOMER (H) database or the JASPAR (J) database. (**B**) The top *de novo* identified motif by HOMER for each subset (STAT1, STAT1/2, STAT2) of the $STAT1_{p701}$ and STAT2 ChIP-seq peak overlaps. The *de novo* motifs were presented as PWM with the probability of each nucleotide on the y-axis and the position within the motif on the x-axis. The log p-value, the percentage of the motifs within the target or in background, best match and similarity scores were calculated with the HOMER *de novo* motif annotation. (**C**) Visualization of the HOMER similarity scores of the Top 10 most similar motifs to the identified *de novo* motifs from **B**. (**D**) Scatter plot of normalized gene expression ($log_{10}$ of TPM) before (0 h, x-axis) and after 6 h (y-axis) of IFNβ stimulation. Genes with $STAT_{p701}$, STAT2 or STAT1/2 binding sites at promoters in ESCs (top) and MEFs (bottom) were shown. Red dots indicate ISGs, while black dots showed STAT-bound genes at the promoter with no significant changes of expression.

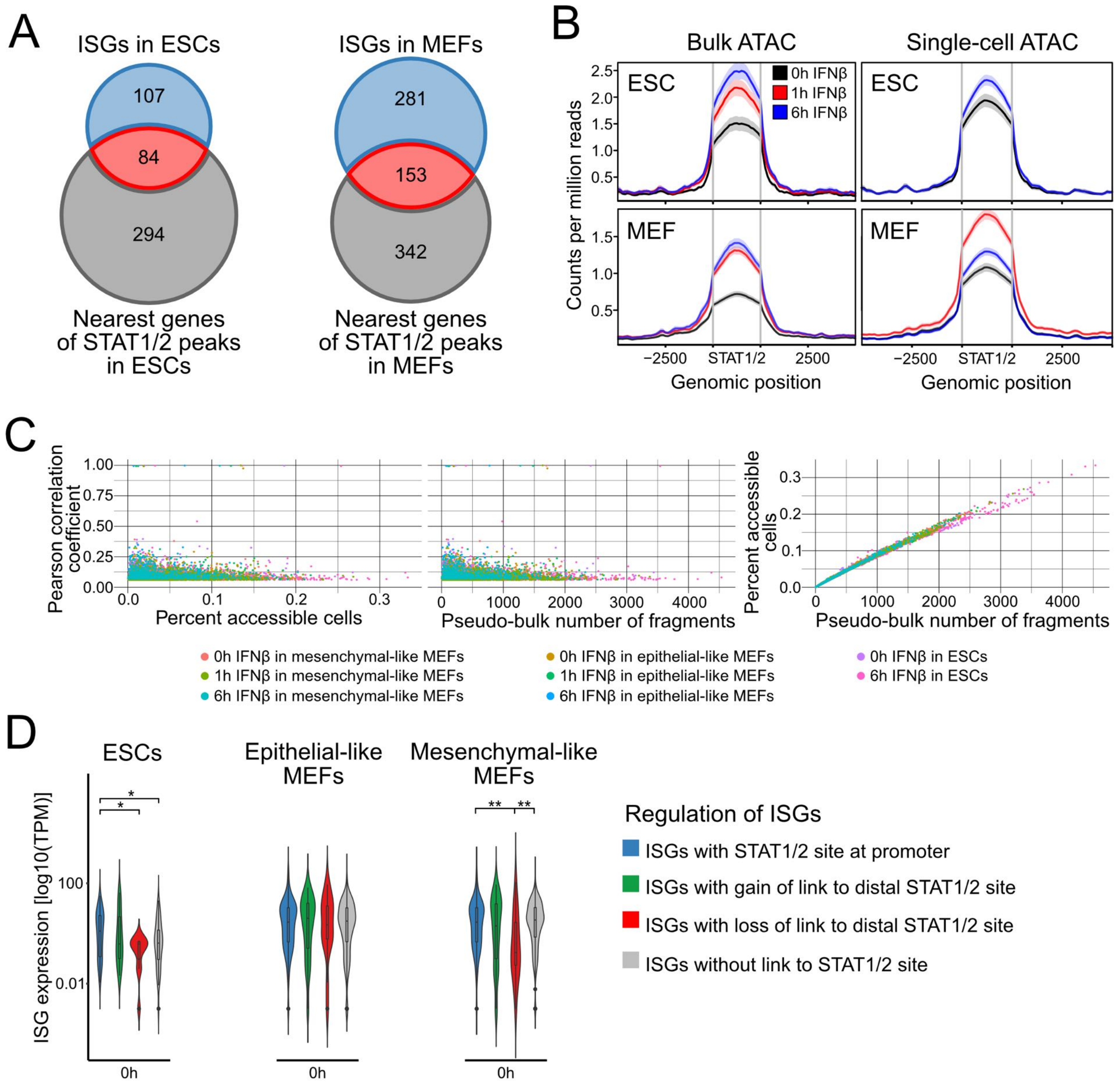


**Supplementary Figure 4. Identification of co-accessible STAT1/2 binding sites and target genes.**

(**A**) Overlap of ISGs identified in ESCs (left) and MEFs (right) with nearest gene to STAT1/2 peak within the corresponding cell type. Nearest gene list was calculated with GREAT and the option "Two nearest genes within [1,000kb]" (McLean et al, 2010). (**B**) Chromatin accessibility at STAT1/2 binding sites in 0 h IFNβ (black), 1 h IFNβ (red) and 6 h IFNβ (blue) ESCs (top) and MEFs (bottom) in bulk (left) and single cell ATAC-seq data (right). A general increase of accessibility at STAT1/2 binding sites upon IFNβ treatment was apparent in single cell and bulk ATAC-seq data except for MEF IFNβ 6h scATAC-seq data. (**C**) Scatter plots of Pearson correlation coefficients, percent accessible cells, and pseudobulk number of fragments for all STAT1/2 co-accessible links above background. The resulting correlation coefficients were independent of general differences in accessibility or more homogeneous accessibility over the cell population in the linked regions. (**D**) Normalized read counts (TPMs) of bulk RNA-seq data for ISGs with different mechanisms of regulation by STAT1/2 binding.

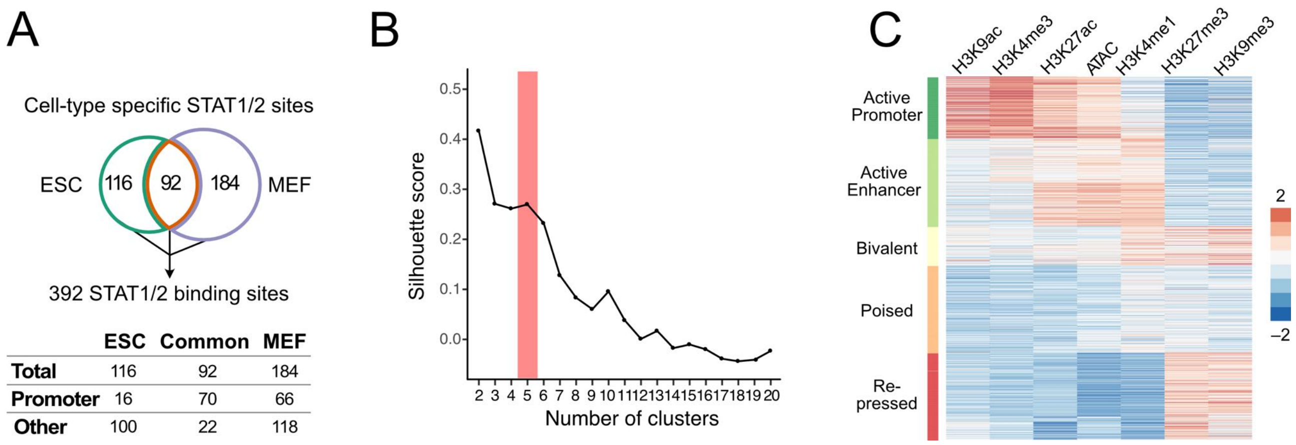


**Supplementary Figure 5. Clustering of chromatin features at STAT1/2 binding sites.**

(**A**) STAT1/2 binding site sites used in the chromatin context analysis. A combined set of 392 STAT1/2 binding sites was obtained from the STAT ChIP-seq analysis in ESCs and MEFs at 1 h and 6 h of IFNβ treatment. Chromatin features in a genomic region of +/-1 kb around the centers of binding sites were analyzed. STAT1/2 binding sites at promoters were annotated according to transcription start sites in the ENSEMBL database. (**B**) Silhouette score calculated as the mean Silhouette coefficient over all samples plotted against cluster number. The clustering of the data described in panel C was evaluated. When varying the cluster number between 2 to 20 clusters it was seen that the selected number of 5 cluster was appropriate for this data set. (**C**) Heatmap with unsupervised k-means clustering of histone modifications (H3K4me1, H3K4me3, H3K27ac, H3K9ac, H3K27me3, H3K9me3) and chromatin accessibility (ATAC) data from unstimulated ESCs and MEFs at 392 STAT1/2 binding sites. Five biologically relevant clusters with distinct signatures were identified and annotated as “Active Promoter”, “Active Enhancer”, “Bivalent”, “Poised” and “Repressed” chromatin states.

**Supplementary Table S1. Different sequencing readouts and replicate number.**

| Readout | Target | Cell type | Treatment | Repli-cates | GEO ID (GSE160764) |
|---|---|---|---|---|---|
| RNA-seq | Total RNA | ESC | 0 h; 1 h; 6h | 4 | GSM4878858 - GSM4878869 |
| | | MEF | 0 h; 1 h; 6h | 2 | GSM4878870 - GSM4878875 |
| | | NPC | 0 h; 1 h; 6h | 4 | GSM4878876 - GSM4878887 |
| scRNA-seq | Poly-A RNA in single cells | ESC | 0 h; 1 h; 6h | 1 | GSM4878890 - GSM4878892 |
| | | MEF | 0 h; 1 h; 6h | 1 | GSM5852363 - GSM5852365 |
| ChIP-seq | STAT1p701 | ESC | 0 h; 1 h; 6h | 4-5 | GSM4878806 - GSM4878819 |
| | | MEF | 0 h; 1 h; 6h | 2 | GSM4878846 - GSM4878851 |
| | STAT2 | ESC | 0 h; 1 h; 6h | 4-5 | GSM4878820 - GSM4878833 |
| | | MEF | 0 h; 1 h; 6h | 2 | GSM4878852 - GSM4878857 |
| | IgG rabbit STAT | ESC | 0 h; 1 h; 6h | 4-5 | GSM4878778 - GSM4878791 |
| | | MEF | 0 h; 1 h; 6h | 2 | GSM4878834 - GSM4878839 |
| | Input STAT | ESC | 0 h; 1 h; 6h | 4-5 | GSM4878792 - GSM4878806 |
| | | MEF | 0 h; 1 h; 6h | 2 | GSM4878840 - GSM4878845 |
| | H3 | ESC | 0 h; 1 h; 6h | 3 | GSM4878706 - GSM4878714 |
| | | MEF | 0h | 3 | GSM5852366 - GSM5852367 |
| | H3K4me1 | ESC | 0 h; 1 h; 6h | 2 | GSM4878730 - GSM4878735 |
| | | MEF | 0h | 2 | GSM5852372 - GSM5852373 |
| | H3K4me3 | ESC | 0 h; 1 h; 6h | 2 | GSM4878736 - GSM4878741 |
| | | MEF | 0h | 2 | GSM5852374 - GSM5852375 |
| | H3K9ac | ESC | 0 h; 1 h; 6h | 3 | GSM4878742 - GSM4878750 |
| | | MEF | 0h | 2 | GSM5852376 - GSM5852377 |
| | H3K9me3 | ESC | 0 h; 1 h; 6h | 3 | GSM4878751 - GSM4878759 |
| | | MEF | 0h | 2 | GSM5852378 - GSM5852379 |
| | H3K27ac | ESC | 0 h; 1 h; 6h | 2 | GSM4878715 - GSM4878720 |
| | | MEF | 0h | 2 | GSM5852368 - GSM5852369 |
| | H3K27me3 | ESC | 0 h; 1 h; 6h | 3 | GSM4878721 - GSM4878729 |
| | | MEF | 0h | 2 | GSM5852370 - GSM5852371 |
| | IgG rabbit histone | ESC | 0 h; 1 h; 6h | 3 | GSM4878760 - GSM4878768 |
| | Input histone | ESC | 0 h; 1 h; 6h | 3 | GSM4878769 - GSM4878777 |
| | | MEF | 0h | 2 | GSM5852380 - GSM5852381 |
| ATAC-seq | Open chromatin | ESC | 0 h; 1 h; 6h | 2 | GSM4878694 - GSM4878699 |
| | | MEF | 0 h; 1 h; 6h | 2 | GSM4878700 - GSM4878705 |
| scATAC-seq | Open chromatin in single cells | ESC | 0 h; 6h | 1 | GSM4878888 - GSM4878889 |
| | | MEF | 0 h; 1 h; 6h | 1 | GSM5852360 - GSM5852362 |

**Supplementary Table S2. Cell type specific ISGs and STAT1/2 binding sites.**

| | | ESC all | MEF all | NPC all | ESC-specific | Common ESC & MEF | MEF-specific |
|---|---|---|---|---|---|---|---|
| **ISGs**[a] | Total | 191 | 463 | 204 | 33 | 158 | 305 |
| | 1 h IFNβ | 57 | 115 | 75 | 18 | 39 | 76 |
| | 6 h IFNβ | 188 | 452 | 240 | 32 | 156 | 296 |
| | Intronic RNA (6h IFNβ) | 82 | 453 | 128 | 8 | 74 | 379 |
| **ChIP-seq peaks**[b] | $STAT1_{p701}$ all | 1,133 | 426 | n. d. | 988 | 145 | 280 |
| | STAT2 all | 236 | 574 | n. d. | 116 | 120 | 453 |
| | $STAT1_{p701}$ only | 925 | 150 | n. d. | 887 | 38 | 112 |
| | STAT2 only | 28 | 298 | n. d. | 25 | 3 | 295 |
| | STAT1/2 | 208 | 276 | n. d. | 116 | 92 | 184 |

[a] ISGs determined from differential RNA-seq analysis. n. d., not determined.

[b] ChIP-seq peaks of $STAT1_{p701}$ and STAT2 called against ChIP-seq of histone H3. STAT1/2 peaks were identified by intersecting the $STAT1_{p701}$ and STAT2 peaks.

**Supplementary Table S3. scATAC-seq data overview and quality.**

| **Cell line** | **ESC** | | **MEF** | | | | | |
|---|---|---|---|---|---|---|---|---|
| Treatment | WT (IFNß0h) | IFNß6h | WT (IFNß0h) | | IFNß1h | | IFNß6h | |
| Cell number | 8,925 | 5,596 | 11,656 | | 12,272 | | 19,403 | |
| Fragments/cell (median) | 16,397 | 24,512 | 12,799 | | 12,095 | | 4,816 | |
| Fraction frag-ments at targeted region | 65.3% | 65.8% | 68.1% | | 64.0% | | 68.5% | |
| | | | Minor cluster | Major cluster | Minor cluster | Major cluster | Minor cluster | Major cluster |
| Sampled cell number | 2,700 | 2,700 | 2,700 | 2,700 | 2,700 | 2,700 | 2,700 | 2,700 |
| Fragments/cell (sampled, median) | 13,443 | 20,143 | 13,452 | 13,454 | 13,463 | 13,460 | 6,557 | 10,544 |

**Supplementary Table S4. Antibodies used in this study.**

| Target | Company | Ref. | Species | ChIP-seq | Western blot |
|---|---|---|---|---|---|
| H3K4me1 | Abcam | ab8895 | Rabbit | 2 µg for 25 µg of chromatin | 1:500 |
| H3K4me3 | Abcam | ab8580 | Rabbit | 2µg for 25µg of chromatin | 1:1000 |
| H3K9ac | Active Motif | 39137 | Rabbit | 10 µl per ChIP | 1:1000 |
| H3K9me3 | Abcam | ab8898 | Rabbit | 2-4 µg for 25 µg of chromatin | – |
| H3K27ac | Abcam | ab4729 | Rabbit | 2 µg for 25 µg of chromatin | 1:1000 |
| H3K27me3 | Abcam | ab6002 | Mouse | 5-10 µg for 25 µg of chromatin | 1:1000 |
| H3K27me3 | Active Motif | 39155 | Rabbit | 5 µg per ChIP | 1:1000 |
| H3 | Abcam | ab1791 | Rabbit | 2µg for $10^6$ cells | 1:1000 |
| IgG rabbit | Acris | AB-105-C | Rabbit | 2 µl | |
| STAT1 | Cell Signaling | #9172 | Rabbit | 1:50 | 1:1000 |
| STAT1 p701 | Cell Signaling | #7649 | Rabbit | 1:100 | 1:1000 |
| STAT1 p727 | Cell Signaling | #8826 | Rabbit | 1:50 | 1:1000 |
| STAT2 | Cell Signaling | #72604 | Rabbit | 1:50 | 1:1000 |
| IgG Rb | Cell Signaling | #2729 | Rabbit | 2 µl (µg/µl) | – |

**Supplementary Table S5. Data analysis software.**

| Software | Reference | Link | Version |
|---|---|---|---|
| ArchR | (Granja et al, 2021) | github.com/GreenleafLab/ArchR | 1.0.1 |
| bedtools | (Quinlan & Hall, 2010) | bedtools.readthedocs.io/en/latest/ | 2.27.1 |
| Bioconductor | (Gentleman et al, 2004) | www.bioconductor.org | 3.10 |
| Bowtie2 | (Langmead & Salzberg, 2012) | bowtie-bio.sourceforge.net/bowtie2/index.shtml | 2.3.3 |
| Cell Ranger scATAC | (Satpathy et al, 2019) | support.10xgenomics.com/single-cell-atac/software/pipelines/latest/what-is-cell-ranger-atac | 1.1.0 |
| Cell Ranger scRNA | (Zheng et al, 2017) | support.10xgenomics.com/single-cell-gene-expression/software/pipelines/latest/what-is-cell-ranger | 3.0.2 |
| DAVID | (Huang da et al, 2009) | david.ncifcrf.gov | 6.8 |
| DESeq2 | (Love et al, 2014) | doi.org/10.18129/B9.bioc.DESeq2 | 1.24.0 |
| DiffBind | (Ross-Innes et al, 2012) | doi.org/10.18129/B9.bioc.DiffBind | 2.12.0 |
| DSS | (Wu et al, 2013) | doi.org/10.18129/B9.bioc.DSS | 2.38.0 |
| Enriched Heatmap | (Gu et al, 2018) | doi.org/10.18129/B9.bioc.EnrichedHeatmap | 3.12 |
| GenomicRanges | (Lawrence et al, 2013) | doi.org/10.18129/B9.bioc.GenomicRanges | 1.36.4 |
| gProfileR | (Reimand et al, 2016) | biit.cs.ut.ee/gprofiler/ | 0.2.0 |
| GREAT | (McLean et al, 2010) | bejerano.stanford.edu/great/public/html/index.php | 4.0.4 |
| HOMER | (Heinz et al, 2010) | homer.ucsd.edu/homer/ | 4.9 |
| HTSeq | (Anders et al, 2015) | htseq.readthedocs.io/en/master/ | 0.12.4 |
| Integrative Genomics Viewer (IGV) | ((Robinson et al, 2011) | software.broadinstitute.org/software/igv/ | 2.3.23 |
| MACS2 | (Feng et al, 2012; Zhang et al, 2008) | github.com/taoliu/MACS | 2.1.2 |
| R software package | (R Core Team, 2020) | www.r-project.org | 3.6.3, 4.0.2 |
| REVIGO | (Supek et al, 2011) | http://revigo.irb.hr/ | n. a. |
| RSEM | (Li & Dewey, 2011) | https://github.com/deweylab/RSEM | 1.3.0 |
| RWire | (Mallm et al, 2019) | https://github.com/FabianErdel/RWire | n. a. |

| | | | |
|---|---|---|---|
| SAMtools | (Li et al, 2009) | http://samtools.sourceforge.net/ | 1.3 |
| SICER | (Xu et al, 2014) | https://github.com/dariober/SICERpy | 0.1.1 |
| SortMeRNA | (Kopylova et al, 2012) | https://bioinfo.lifl.fr/RNA/sortmerna/ | 2.1 |
| STAR | (Dobin et al, 2013) | github.com/alexdobin/STAR | 2.5.3a |
| Trimmomatic | (Bolger et al, 2014) | www.usadellab.org/cms/?page=trimmomatic | 0.36 |
| Seurat | (Stuart et al, 2019) | satijalab.org/seurat/ | 4.0.1 |
| Venny | n. a. | bioinfogp.cnb.csic.es/tools/venny | 2.1 |

**Supplementary Table S6. Inventory of Supplementary Data Sets.**

| File Name | Figure/ table ref. | Description |
|---|---|---|
| Supplementary Data Set 1: Dataset_01_ISG.xlsx | Fig. 1, 2; Fig. S1, S2; Table S2; | ISGs identified in ESCs, NPCs and MEFs after 1 h and 6 h treatment. Assignment of cell type specific and common ISGs in ESCs and MEFs. |
| Supplementary Data Set 2: Dataset_02_STAT.xlsx | Fig. 3; Fig. S3; Table S3; | Binding sites of STAT 1 and STAT 2 (only STAT1 or 2 as well as common binding sites) in ESCs and MEFs. Binding site motifs from anlaysis of known or de nove identified motifs. |
| Supplementary Data Set 3: Dataset_03_ISG-regulation.xlsx | Fig. 4; Fig. S3, S4; Table S3; | Assignment of regulatory mechanism to ISGs, i.e. promoter binding of STAT1 and/or STAT2 as well as assignment of STAT1/2 bound enhancers predicted from the co-regulation analysis. |
